# Novel phages of *Pseudomonas syringae* unveil numerous potential auxiliary metabolic genes

**DOI:** 10.1101/2024.05.07.591244

**Authors:** Chloé Feltin, Julian R. Garneau, Cindy E. Morris, Annette Bérard, Clara Torres-Barceló

**Author notes:** **Repositories:** Assembled and annotated genomes of the 25 *Pseudomonas syringae* phages were uploaded to GenBank under accession numbers PP179310 to PP179334. All data generated or analysed during this study are included in this published article (and in the two “Supplementary Material” files).

## Abstract

Relatively few phages that infect plant pathogens have been isolated and investigated. The *Pseudomonas syringae* species complex is present in various environments, including plants. It can cause major crop diseases, such as bacterial canker on apricot trees. This study presents a collection of 25 unique phages genomes that infect *P. syringae*. These phages were isolated from apricot orchards with bacterial canker symptoms after enrichment with 21 strains of *P. syringae*. This collection comprises mostly virulent phages, with only three being temperate. They belong to 14 genera, 11 of which are newly discovered, and 18 new species, revealing great genetic diversity within this collection. Novel DNA packaging systems have been identified bioinformatically in one of the new phage species, but experimental confirmation is required to define the precise mechanism. Additionally, many phage genomes contain numerous potential auxiliary metabolic genes with diversified putative functions. At least three phages encode genes involved in bacterial tellurite resistance, a toxic metalloid. This suggests that viruses could play a role in bacterial stress tolerance. This research emphasises the significance of continuing the search for new phages in the agricultural ecosystem to unravel novel ecological diversity and new gene functions. This work contributes to the foundation for future fundamental and applied research on phages infecting phytopathogenic bacteria.

## 3. Introduction

As the most abundant biological entities on the planet, the viruses of bacteria or bacteriophages (phages), play a major role in diverse ecosystems. Knowledge of these roles has provided better insight into the dynamics of diverse bacterial communities including those that govern the major biogeochemical cycles in the marine environment [1, 2], in soil [3], and in the human gut [4].

According to several studies, it has been established that phages can exert a significant influence, both quantitatively and in terms of diversification, on the host bacterial population [5, 6]. When phages are studied with respect to agriculture, they are often considered as biocontrol agents against plant pathogenic bacteria (for instance *Xanthomonas oryzae* pv*. oryzae*: [7]; *Ralstonia solanacearum:* [8]; *Pseudomonas syringae* pv*. actinidiae*: [9]; *Pectobacterium spp.*[10]). The use of phages against bacterial diseases in agriculture was proposed very early on after their discovery [11–14]. Today’s world challenges are driving the need for more sustainable agricultural solutions, such as phage-based products, which are an example of biocontrol strategy. To develop these environmentally-friendly alternative solutions and better understand their interactions within microbial communities, it is necessary to identify and characterise phages.

Among the plant pathogenic bacteria that cause significant damage in agriculture, *P. syringae* is a species complex, classified into at least 13 phylogroups (PG) and 23 clades based on multilocus sequencing typing (MLST) [15]. *P. syringae* is ubiquitous and associated with ecosystems linked to the water cycle, such as lakes, rivers, clouds, snow, biofilms, soil litter, wild and cultivated plants [16]. In the latter case, this bacterium can cause losses of up to 60% in arboriculture and horticulture [17]. This pathogen can colonise leaves and other aerial plant parts as an epiphyte from which it can penetrate natural openings and wounds to attain the intercellular spaces as an endophyte. In some cases, inoculum from the endophytic phase can be devastating for the plant with sudden wilting, the presence of cankers on stems and main trunks, and the production of exudates [18]. On apricots, *P. syringae* can cause bacterial cankers which lead to a complex and episodic disease with considerable economic impacts in several regions of the world, particularly in Iran [19], Turkey [20], Bulgaria [21] and Italy [22]. Controlling bacterial canker on apricots is difficult due to the complex nature of the disease and the responsible pathogen. The *P. syringae* strains responsible for the disease represent a high degree of genetic variability [23, 24]. On apricot tissues affected by bacterial canker, the pathogenic strains are mainly identified as PG01, PG02, and less frequently as PG03 and PG07 [23].

The pathogenicity of a strain on apricot trees cannot be predicted either by its phylogeny or by phenotypic characteristics, and phages may be playing an important, yet unknown, role.

Phages infecting human pathogenic bacteria have been studied more extensively than those infecting phytopathogenic bacteria. This can be measured by the number of phage genomes sequenced: 2017 accessions for *Escherichia coli* phages and 873 for *P. aeruginosa* phages, whereas there are only 110 *P. syringae* phage genomes (INPHARED; March 2024). The diversity of the *P. syringae* species complex is comparable to the bacterial complexes of these two other families, indicating that further diversity is yet to be discovered for *P. syringae* phages. The need to expand phage genome databases therefore remains important, as does the need to build up collections of purified phages relevant to agriculture contexts.

This imbalance is changing, as illustrated by the growing literature on phages of plant-associated bacteria. Recent studies on *P. syringae* phages have investigated their potential as biocontrol agents. For instance, the Medea1 phage represented a new genus and its biocontrol capacity was tested *in vitro* and *in planta* [25]. It was isolated from a strain of *P. syringae* pv. *tomato* from tomato seedling soil and characterised genetically and phenotypically. This new phage also showed strong antibiofilm activity. Similarly, the phage ΦPsa374 was isolated from soil and compost by enrichment with *P. syringae* pv. *actinidae* strains [9]. Their study described the ΦPsa374 genome to understand its properties and infection strategies, with the aim of using it as a durable antimicrobial agent. Warring *et al.* showed that this phage uses LPS as a receptor *in vitro* and *in planta*, and that mutations in the synthesis of these LPS are involved in resistance to phage infection [26]. Numerous mutations in the phage genome were also noted in the tail fiber and a structural protein, suggesting evolution and adaptation of the phage to the bacterial strains. ΦPsa374 genome sequencing included genes associated with tellurium resistance, a rare trait that has been reported to possess antimicrobial properties. These types of genes are auxiliary metabolic genes (AMGs) integrated into the phage genome and are likely to promote the fitness of the infected bacteria and thus indirectly enhance phage replication efficiency. Although these genes have been described in other ecological contexts (marine [27] or soil [28]), to our knowledge, no studies has described AMGs in phages of plant-associated bacteria. These recent findings highlight the potential for finding phages particularly suited for biocontrol, as well as the prospect of discovering novel AMGs.

In this work, we sought to collect of phages from an agricultural context where *P. syringae* is an important pathogen, viz. as the causal agent of bacterial canker of apricots. The soil of symptomatic apricot orchards was sampled and enriched with several *P. syringae* strains with the objective of detecting a wide diversity of phages. The genetic description of this phage collection was accompanied by functional genome annotation and inter-genus comparison to identify interesting characteristics of newly discovered phage genera. This new collection highlights the important genetic diversity found in *P. syringae* phages.

## 4. Methods

### 4.1.1 Soil sampling in apricot orchards

All samples were taken in November 2021 at the INRAE experimental research unit in Gotheron (Saint-Marcel-lès-Valence, Drôme, France) in three apricot orchards. Supplementary table 1 describes the characteristics of these orchards. Ten soil samples were taken from each orchard. Soil was sampled from the base of the tree showing symptoms of bacterial canker (within a perimeter of approximately 30 cm at a depth of approximately 10 cm (one 50 ml Falcon tube per sample from the first layer of soil or the A horizon). The soils had a cation exchange capacity of 12.2 meq/100g, a neutral pH (pH water 7.0) and an organic matter level of 1.9%.

### 4.1.2 Selection of bacterial strains and culture conditions

Twenty-one strains of *P. syringae* from the Plant Pathology Research Unit (INRAE, Montfavet, France) collection were used for phage isolation (table 1). It should be noted that those bacterial strains were not isolated during the course of that study. These strains reflect the genetic diversity of *P. syringae* on apricot (table 1) [23]. It also included *P. syringae* strains isolated from other host plants and from aquatic environments genetically close to strains isolated on apricot on the basis of the sequence of the housekeeping gene citrate synthase (*cts*) (table 1). Phylogenetic analyses were conducted using MEGA version 11 [29] by building Neighbour-Joining tree with 1 000 bootstraps using the p-distance method and rooted with strains outside of the species complex. All the strains belong to five different phylogroups (PG01, PG02, PG03, PG07, PG10) with at least 3 strains per phylogroup except for PG10 (table 1). *P. syringae* strains were stored at -80°C in phosphate buffer with glycerol (20% vol/vol). They were cultured on King’s B (KB) agar medium [30] for 48 hours at 25°C before each experiment.

**Table 1:** The 21 strains of *P. syringae* used for the phage isolation enrichment step. The isolation characteristics of *P. syringae* strains are presented as well as phylogenetic and phenotypic traits.

| Environment of origin | Names | CFBP collection | Phylogroup | Pathogenicity<sup>a</sup> | Country of isolation | Date of isolation | Plant and environment of isolation | Number of phage isolates |
| --- | --- | --- | --- | --- | --- | --- | --- | --- |
| Haplotypes isolated from apricot trees | DG1a.21 |  | PG01a | + | France | 13/04/2015 | *Prunus armeniaca* | 5 |
|  | P1.01.01B03 |  | PG01a | - | France | 01/03/2014 | *Prunus armeniaca* | 14 |
|  | 19B |  | PG01a | + | France | 16/03/2011 | *Prunus armeniaca* | 4 |
|  | P3.01.09C09 |  | PG01a | - | France | 01/03/2014 | *Prunus armeniaca* | 15 |
|  | 41A |  | PG02b | + | France | 14/04/2011 | *Prunus armeniaca* | 0 |
|  | P4.01.01C03 |  | PG02b | - | France | 01/05/2014 | *Prunus armeniaca* | 9 |
|  | DG12.6 |  | PG02d | + | France | 13/04/2015 | *Prunus armeniaca* | 17 |
|  | 7C |  | PG02d | + | France | 16/03/2011 | *Prunus armeniaca* | 22 |
|  | DG8.15 |  | PG07 | + | France | 13/04/2015 | *Prunus armeniaca* | 7 |
|  | 3A |  | PG07 | - | France | 16/03/2011 | *Prunus armeniaca* | 4 |
|  | 93D |  | PG07 | + | France | 14/04/2011 | *Prunus armeniaca* | 10 |
|  | 93A |  | PG03 | + | France | 14/04/2011 | *Prunus armeniaca* | 9 |
| Haplotypes isolated from other plants | CC1630 | CFBP 8561 | PG01a | nt | USA | 27/10/2007 | *Onobrychis viciifolia* | 0 |
|  | DC3000 | CFBP 7438 | PG01a | nt | United Kingdom | 01/01/1960 | *Solanum lycopersicum* | 1 |
|  | CC0440 | CFBP 8573 | PG02b | nt | France | 13/06/2002 | *Cucumis melo* | 4 |
|  | CC0094 | CFBP 5467 | PG02d | nt | France | 22/06/1997 | *Clematis patens* | 18 |
|  | CC0663 |  | PG07 | nt | France | 20/04/2004 | *Primula officinalis* | 23 |
|  | JD03 | CFBP 8535 | PG03 | nt | USA | 01/01/1977 | *Glycine max* | 9 |
|  | Pph1448A |  | PG03 | nt | Ethiopia | 01/01/1985 | *Phaseolus vulgaris* | 13 |
| Haplotypes isolated from water environments | USA0011 | CFBP 8553 | PG02d | nt | USA | 12/08/2007 | River | 16 |
|  | CC1557 | CFBP 8533 | PG10b | nt | France | 20/06/2006 | Rain | 23 |

If present in the CFBP collection, the strain's reference number in that collection is provided. The CIRM-CFBP is the French collection for plant-associated bacteria (https://cirm-cfbp.fr/ ; Feb 2024). The pathogenicity of the strains on apricot tree is detailed as described by Parisi *et al.* (2019) : nt = not tested; + = pathogenic; - = not pathogenic

### 4.1.3 Phage isolation

The 30 soil samples were stored at 4°C in 50 ml Falcon tubes for 3 weeks. Isolation was carried out according to standard protocols [31]. Briefly, the homogenisation phase consisted of mixing approximately 50 g of soil with 110 ml of liquid KB medium with slow agitation for 1 hour at 25°C. The supernatant was centrifuged (3215 ***g*** for 5 min) and then filtered (Merck Millex™ Syringe Filter, PES, 0.22 μm). In the enrichment phase, this filtrate was mixed with each of the 21 *P. syringae* strains (table 1) in exponential phase (OD= 0.2 corresponding to approximately 10^8^ c.f.u. ml^-1^). This mixture was incubated at 25°C for 48 h with slow agitation. Only the phages were kept by adding 10% chloroform, then vortexing and centrifuging (16,873 ***g*** for 5 min) to retain the supernatant. To observe the presence of phage in these enrichments, screening was carried out with the “spot test” technique using the double-layer agar method [32]. From the 221 positive screening tests showing halos of bacterial lysis, 89 were selected for phage purification, representing a variety of strains of *P. syringae* and soil samples. To purify phages, p.f.u. (bacterial lysis plaque-forming units) were isolated three consecutive times using the double-layer agar method. A total of 50 phages (56%) were isolated and purified in 100 µl of SM buffer (100 mM NaCl, 8 mM, Mg SO4 7H2O, 50 mM Tris-HCl, pH 7.4). Finally, phages were amplified at a titre ranging from 10^4^ p.f.u. ml^-1^ to 10^11^ p.f.u. ml^-1^. The selection of reference species phages was based on those with a high p.f.u. ml^-1^ titre that remain stable over time. The phage collection was stored at 4°C in SM buffer and at -80°C in glycerol (50% vol/vol). In addition, attempts were made to isolate phages from the soil of asymptomatic trees, but the lysis plaques were too turbid to purify the phages. Therefore, no phages were purified from the soil of asymptomatic trees.

### 4.1.4 Phage morphology observed by TEM

Electron micrographs of *P. syringae* phages were generated as previously described [33]. Briefly, the sample was placed on a 400-mesh copper grids (Delta microscopies, Mauressac, French), negative stained with 1% (W/V) ammonium molybdate, and analysed using a HT7800 Hitachi transmission electron microscope (Hitachi, Tokyo, Japan) at an acceleration voltage of 80 kV. Electron micrographs were taken with an AMT XR401, sCMOS-camera (AMT imaging, Woburn, MA-US).

### 4.1.5 DNA extraction

DNA was extracted from the 50 phages in the collection using either the Phage DNA Isolation kit (Norgen Biotek CORP) or a phenol-chloroform protocol from pure amplified phages. The quantity and quality of the extracted DNA was checked with a nanodrop spectrophotometer (ND-1000 Spectrophotometer), Qbit (3.0 Fluometer) and the DNA was visualised on a gel (1% agarose). The DNA of the 50 phages was then sent to the Biomics sequencing platform at the Pasteur Institute (Paris, France). The libraries were prepared using Illumina® TruSeq PCR-Free and sequencing was carried out using ISeq100 PE150 (2 × 150 pb read length, target 100X / sample). Of the 50 genomes, 38 were successfully sequenced and assembled.

### 4.1.6 Phage genome assembly and annotation

Sequence analysis was performed on Galaxy CPT [34] and genome assembly was performed *de novo* with the Trimmomatic [35], SPAdes [36], PhageTerm [37] tools. Phage morphology was determined with VIRFAM [38]. The phage life cycle was identified with PhageAI (Supp table 3) [39]. The comparison of genomes by inter-genomic distances allowed us to analyse the taxonomy of our collection of 38 phages sequenced relative to close reference phages currently known (VIRIDIC) [40]. The functional annotation of phages was updated using PROKKA [41] with a priority annotation using a modified PHASTER protein database (version Dec 22; [42]). To increase the number of identifiable ORF, hypothetical proteins were removed, and the E-value threshold for function assignment was set to 0.01. The predicted function of all genes was manually categorised (Supp table 4), which also led to the identification of potential AMGs. These potential AMGs were verified by comparing the protein sizes present in the phage and host genomes and by homologous comparison with a strict identity threshold of 70% and coverage of 90%. The INPHARED database was used to count the number of phages infecting *P. syringae* [43].

## 5. Results and Discussion

### 5.1.1 Phages were found in all apricot soils when screened against 21 strains of *P. syringae*

To increase the likelihood of isolating a diverse range of phages, 21 strains of *P. syringae*, representing a wide range of diversity, were each added to all sampled soil (table 1). For the total of 630 enrichments that were screened (30 soil samples x 21 host strains), 221 (35%) yielded a phage isolate and 100% of the soils yielded *P. syringae* phage isolates (*i.e.* at least one phage isolate in each soil sample could attack one of the *P. syringae* strains used for enrichment). An average of 7 phage isolates were counted per soil, regardless of the orchard sampled. Moreover, the number of phages isolated was similar between the 3 orchards, with 76 in the Hybrid orchard (34%), 78 in the CapRED orchard (35%) and 67 in the Core Collection orchard (30%), revealing that there was no effect of orchard on the number of phage isolates (Supp figure 1). This result is not surprising given that the orchards sampled are very close geographically and therefore share climatic and soil characteristics, differing only in the genotype of the apricot trees. Our results prove that the isolation of *P. syringae* phages from soil is efficient. Indeed, it has been shown that *P. syringae* phages are easily isolated from soils and from water samples [44, 45]. A similar study sampled 70 isolates of *P. syringae* phages from 60 cherry trees (soil, leaves and bark) in 6 different geographical locations [46]. Their results were similar to ours since all soil samples yielded phages of *P. syringae*. However, in contrast with our results, the distribution of the number of phages isolated by geographical site was not uniform (45%, 26%, 16%, 7%, 4% and 2%). This result probably depends on the sampling design and accounts for the diversity of climate, soil, type of farm, number of enrichment strains, and geographical distance between sites.

**Figure 1:**
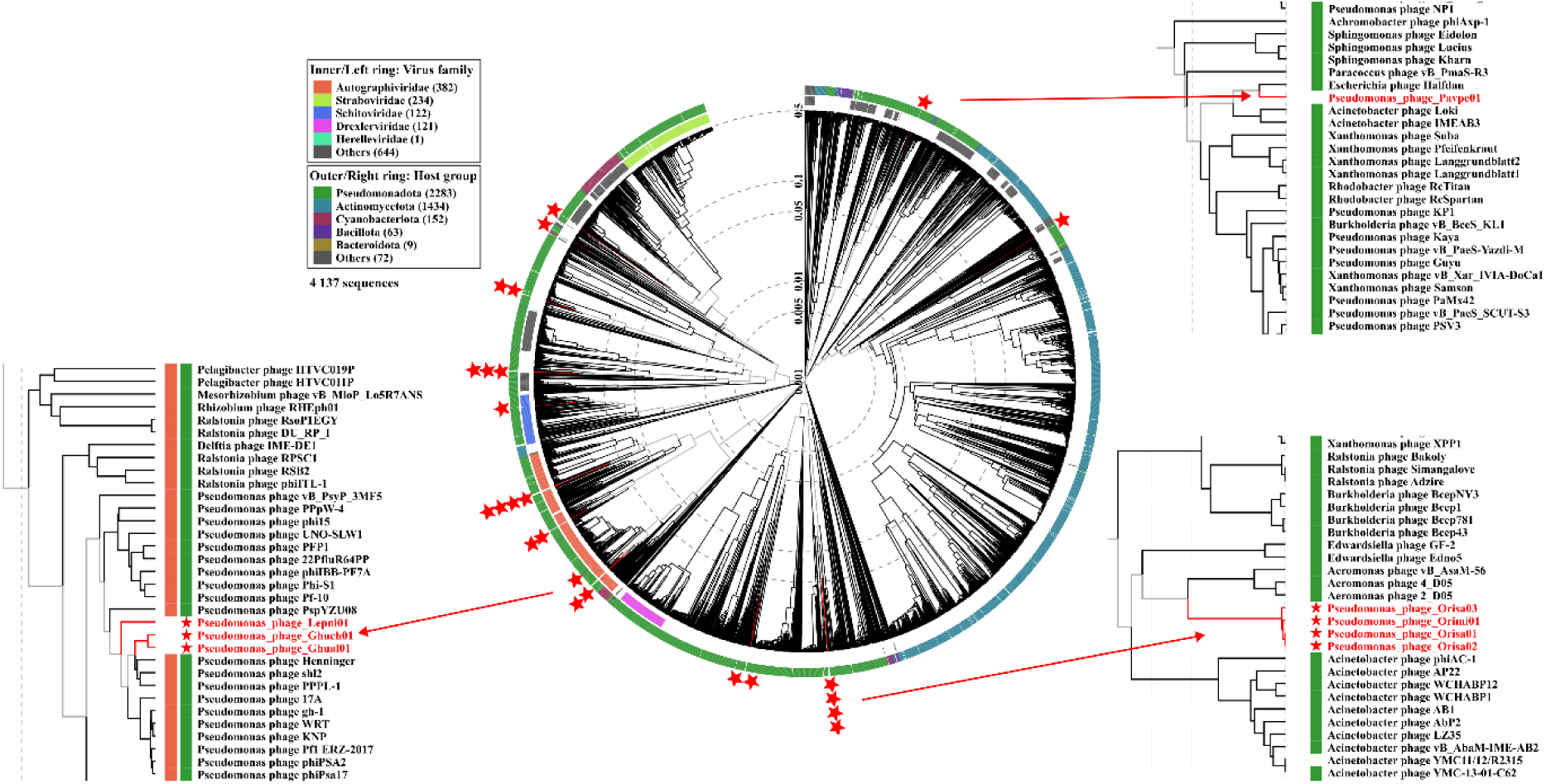
Proteomic tree of the phylogenetic context of 25 newly discovered *P. syringae* phage genomes (indicated by red stars), generated with VipTree and modified with Inkscape version 1.3.2. The inner ring represents the known virus families of the related phage genomes, while the outer ring shows the host group of these phages. Enlarging three groups highlighted some phages close to those in the collection. Phages are indicated with their respective GenBank accessions identifiers.

The relatively wide diversity of *P. syringae* strains used to enrich the phages in the 30 soils sampled from three apricot orchards made it possible to isolate a large diversity of phages (table 1). Of the 21 strains of *P. syringae* used, 19 were host to at least one phage (table 1). On average, each bacterial strain was able to isolate 11 phages from the three orchards. Strain JD03 (PG03) was the most efficient at isolating phages, having isolated 23 phages from the three orchards. Strain 7C (PG02b) was also highly efficient, allowing the isolation of 22 phages (table 1). In contrast, strains CC1630 (PG01a) and 41A (PG02b) did not yield any phage (table 1). Parisi *et al*. tested 7C and 41A for their ability to cause disease on apricots, as well as beans, melons, tomatoes, and cherries [23]. As they have the same host range on the plants tested, pathogenicity does not appear to be linked with their ability to isolate phages. There is no evidence to suggest that a strain of *P. syringae* will be effective in isolating phages according to the habitat (e.g. plants *vs* water) from which they were isolated (table 1). This reflects the fact that the *P. syringae* complex as a whole shows no biogeographical specificity and populations have been mixed among the various substrates in which it can be found [47]. In this work, we find no differences in phage isolation between pathogenic and non-pathogenic strains on apricots. This result was expected since the diversity of *P. syringae* on apricots mainly comprises non-pathogenic strains [23].

Bacteria from all phylogroups were able to isolate phages, with PG02 isolating the highest percentage (38%) followed by PG07 (19%), PG03 (19%), PG01 (18%), and PG10 (7%) (table 1, Supp figure 1). In addition, there was no bacterial phylogenetic pattern that influenced the number of phages isolated. These results justify our approach and our choice of *P. syringae* strains, as they almost all contributed to the diversity of phages collected. The genetic diversity of populations of *P. syringae* on apricot trees is large and represents four phylogroups of the species complex [23].

Phages were isolated from all of them, indicating their ubiquitous presence in the soil of apricot trees. Even if *P. syringae* strains tend to have low residence time in soil [48], rainwater run-off could potentially create a temporary reservoir for them and hence, for their associated phages.

### 5.1.2 *P. syringae* phages from apricot soils represent 11 new genera and 18 new species

A taxonomic analysis of the pure phages indicated a significant amount of novelty and diversity among the phages collected. Calculation of the genetic identity between the 38 purified phages indicated that 66% were unique clones representing 25 unique *P. syringae* phages. The phages belong to 18 new species and 14 genera, 11 of which are new according to ICTV criteria [49] (Supp figure 2), testifying to the great taxonomic diversity of this new collection (table 2). All of the phages in the collection belong to the class of *Caudoviricetes* (Phylum: *Uroviricota*, Kingdom: *Heunggongvirae*, Realm: *Duplodnaviria*) and therefore have the three best-known morphologies: podovirus (52%), myovirus (32%) and siphovirus (16%) (Supp figure 3). The 11 new genera have been named after constellations and the 18 new species have been named after the stars that make up the constellations (table 2).

**Figure 2.**
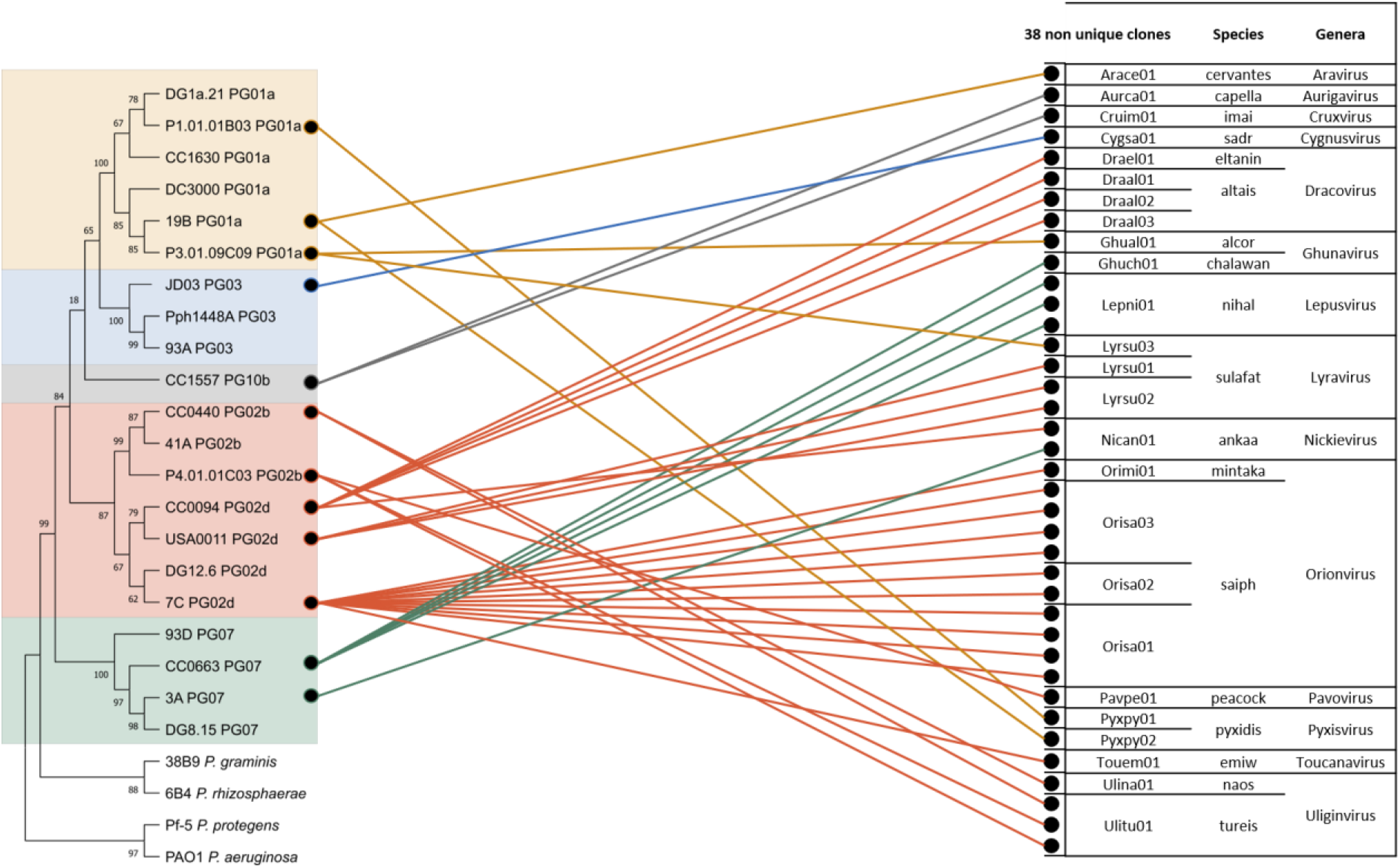
: Network between the enrichment strains of *P. syringae* on the left and the 38 non-unique *P. syringae* phages on the right. The *P. syringae* strains are represented in a Neighbour-Joining tree from the housekeeping cts (citrate synthase) gene and rooted with strains outside of the species complex. The four phylogroups in this tree are delimited by different coloured backgrounds (yellow: PG01a, blue: PG03, pink: PG02b and d, grey: PG10b, green: PG07). The phage collection includes 38 non-unique phages, identified by a dot and named after their reference phage clone. Phages are classified according to their species and genus.

**Figure 3:**
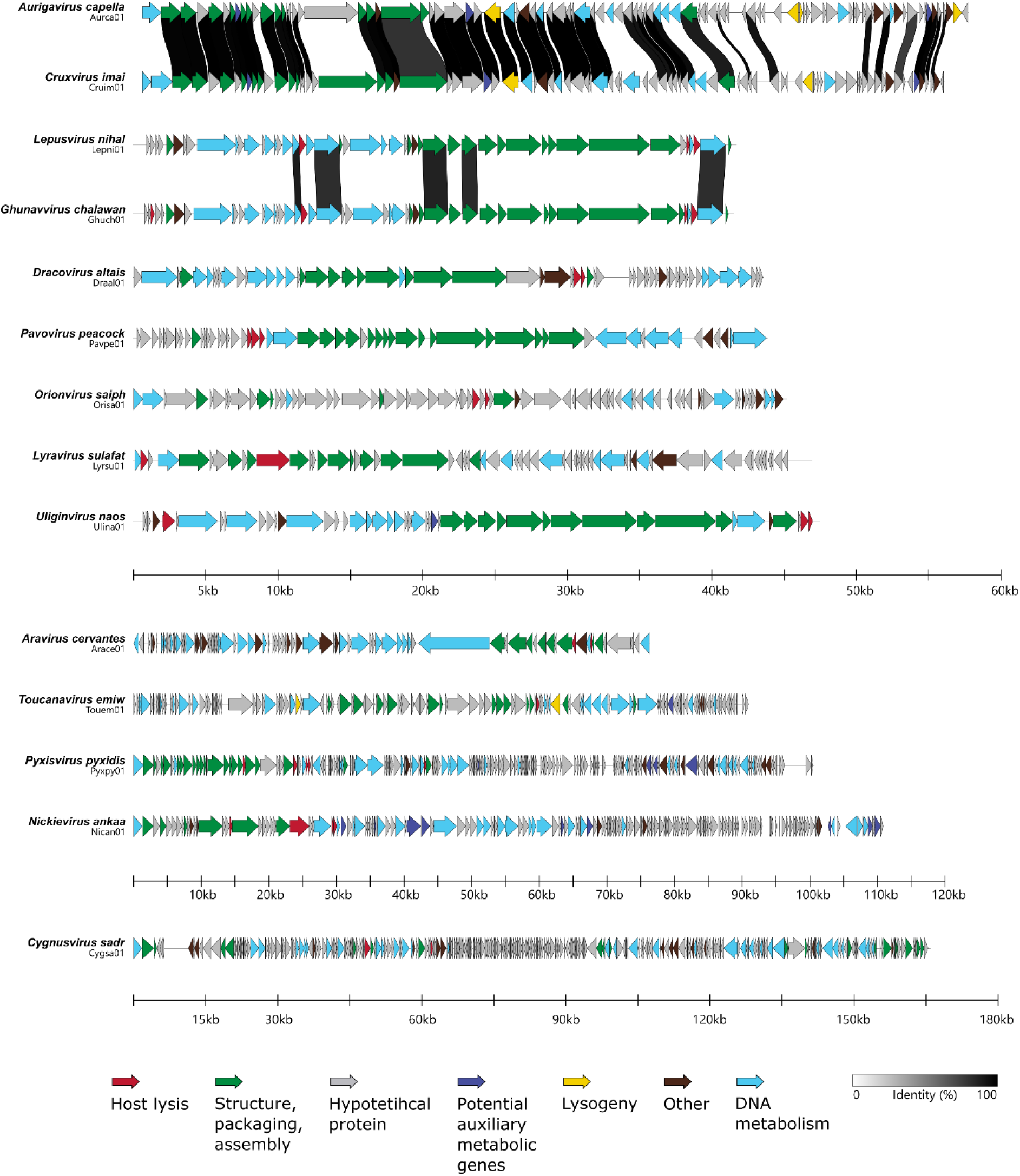
Comparison of the structures of 14 genomes representing the 14 *P. syringae* genera made with Clinker and modified with Inkscape version 1.3.2. The arrows indicate the open reading frame and the direction. Functional modules are indicated and categorised by coloured arrows. Links for shared identity between proteins are presented as black to white bands and protein identity lower than 80% is not shown.

**Table 2:**
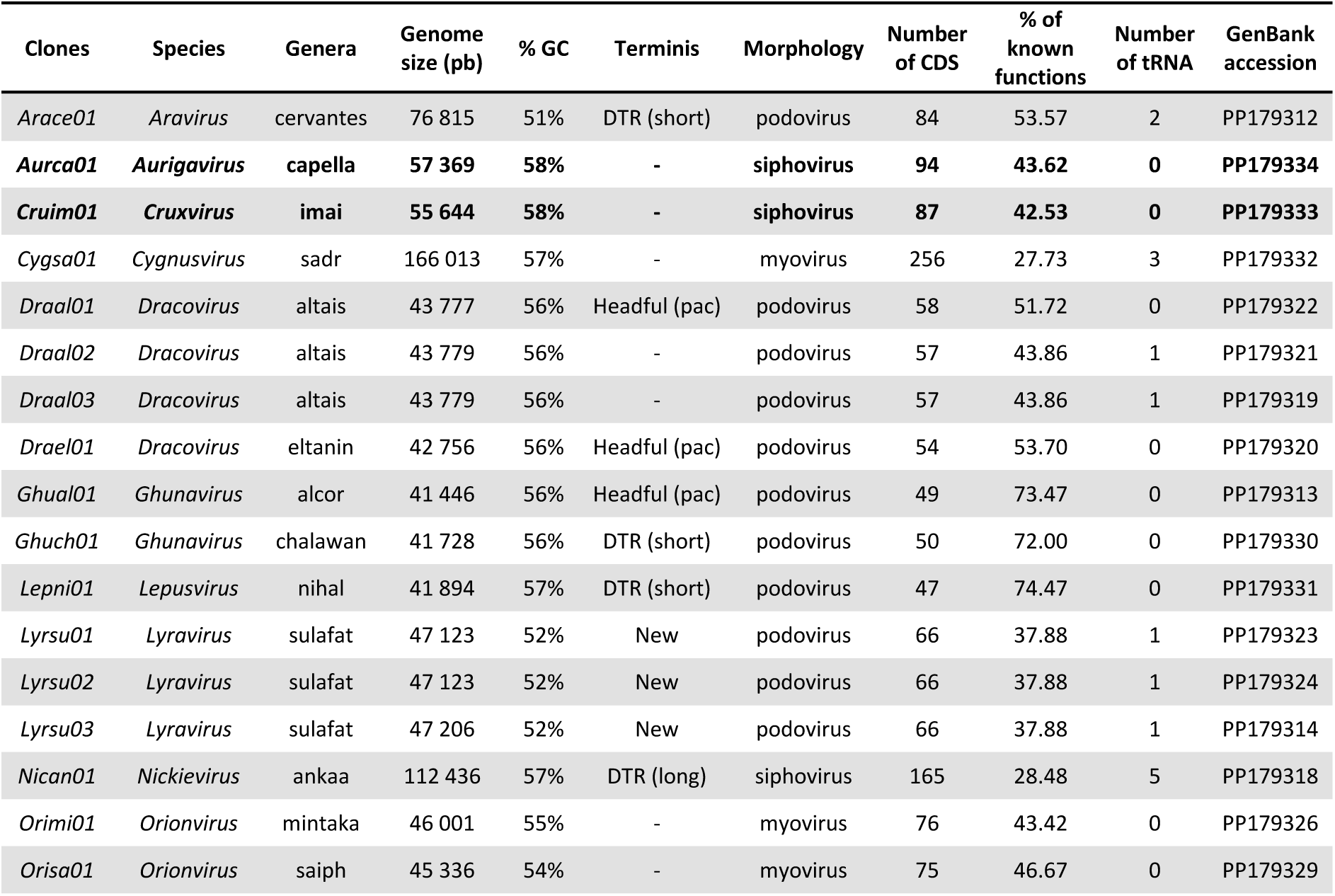
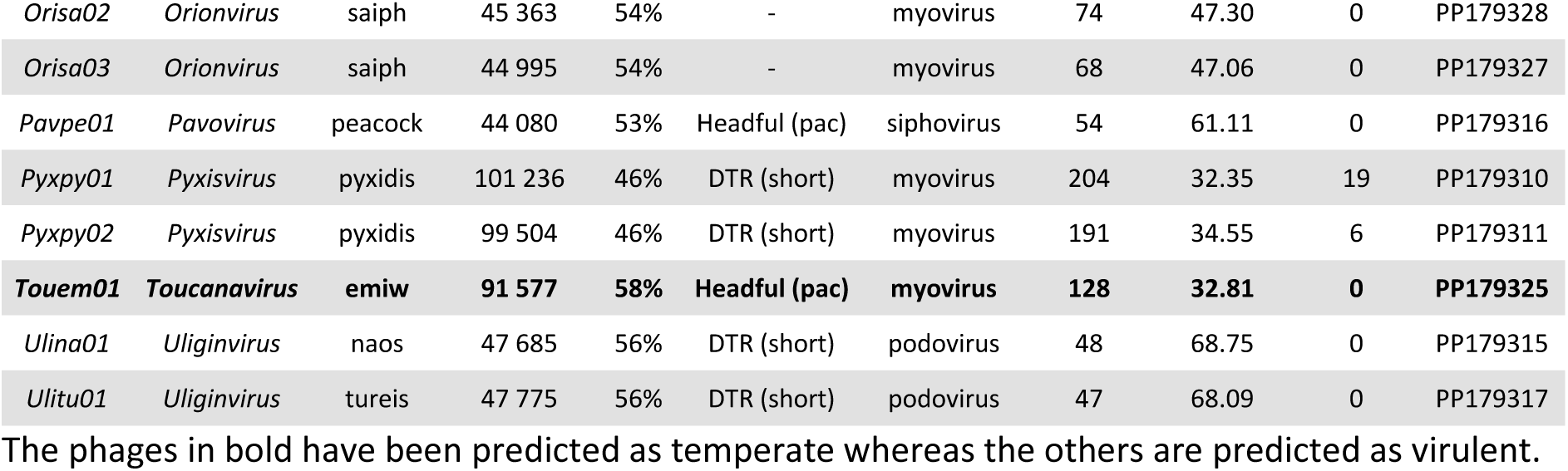
Genetic characteristics of 25 *P. syringae* phages.

To estimate the sampling effort for this study, we compared the number of enrichment strains and the sites sampled to those of a similar study. In their study, Rabiey *et al.* revealed that the 13 purified phages belonged to 5 different genera [46]. They were obtained from 6 cherry orchards located in 6 different geographical sites using 3 enrichment strains. Rabiey’s approach isolated 5 phage genera for 3 enrichment bacterial strains (5/3=1.67), while our approach yielded 14 genera for 21 enrichment strains (14/21=0.67). However, the majority of our diversity was obtained at a single site whereas 6 different sites with 10 varieties sampled were required to capture the diversity obtained by Rabiey *et al*. Considering only the intrinsic diversity of the two collections, ours contains 14 genera for 25 unique phages (25/14=1.79), compared to 5 genera for 13 unique phages (13/5=2.60). As a result, our collection has greater intrinsic diversity due to the lower number of phages per genus and the higher number of genera. For our study, we hypothesised that a high diversity of *P. syringae* phages would require a large number of different enrichment strains compared to other studies that typically used only one or two bacterial strains [25, 26, 50]. Yet, to increase the diversity of *P. syringae* phages with an efficient sampling effort, it seems that using at least 3 enrichment strains (instead of 21) from a single geographical site would enable the collection of many distinct genera of *P. syringae* phages. However, additional phage sampling and enrichment projects should be conducted at contrasted sites to confirm this hypothesis.

The taxonomic diversity of our collection of 25 *P. syringae* phages was analysed at the genus level. There was no inter-genomic similarity between the 14 genera, except for the *Lepusvirus* genus phage, which shared around 58% inter-genetic similarity with the phage belonging to the *Ghunavirus* genus, and the *Aurigavirus* genus phage, which shared 69% with the *Cruxvirus* genus phage (see Supp figure 2). This differentiation between almost all genera was further highlighted by comparing our collection with existing phages in databases, as shown in figure 1. The phages in the collection were widely distributed in the proteomic tree and the closest phages related to ours originated from various hosts, such as *Pseudomonas, Escherichia, Acinetobacter, Bordetella, Aeromonas, Ralstonia*, and *Erwinia* (figure 1). Out of the 11 new genera, 6 are represented by only one phage clone. Only 3 of these 6 genera are composed of temperate phages, indicating a misrepresentation of these *P. syringae* phages. The genera *Dracovirus* and *Orionvirus* are unique in that the isolation strain is common to all four clones of phages within those two genera (CC0094 for *Dracovirus* and 7C for *Orionvirus*, both from phylogroup 02d).

Figure 2 displays the contribution of each enrichment strain to the phage collection for the 38 non-unique phages thereby illustrating how different *P. syringae* strains lead to isolation of identical phage clones. The *Uliginvirus tursei* Ulitu01 phage demonstrates this phenomenon with P4.01.01C03 (PG02b) as the isolation strain and a 100% identical phage clone isolated from the CC0440 (PG02b) enrichment strain. Similarly, the *Nickievirus ankaa* Nican01 phage was isolated from enrichment strain CC0094 (PG02d) and a 100% identical phage was isolated from strain 3A (PG07). The *P. syringae* strains that isolated those phage clones are from completely different phylogroups and were isolated from different plant species. The multiplicity of enrichment strains enabled us to postulate that those strains will be included in the host range of the two unique phages presented below.

### 5.1.3 *P. syringae* phages exhibit a range of genomic size and potential new termini

The size of the genomes and their relationship with existing phages in the GenBank database are the first characteristics that distinguish the different genera in our phage collection. The median genome size was 47.123 kb (*Lyravirus sulafat*), ranging from 41.446 kb (*Ghunavirus alcor*) for the smallest to 166.013 kb (*Cygnusvirus sadr*) for the largest (see table 2). Our collection of phages had an average GC percentage of 54%, with *Pyxisvirus pyxidis* having the lowest at 46% and *Toucanavirus emiw, Cruxvirus imai* and *Aurigavirus capella* having the highest (58%). Furthermore, the GC content of the three temperate phages is the most similar to that of their host *P. syringae*, averaging at 59% across 58 complete genomes (GenBank; Jan 2024). It is worth noting that temperate phages often have an GC content that is more similar to that of their host than virulent phages [51].

A comparison of the phages in the collection with the 15 most closely related phages in the GenBank database is presented in the Supp table 2. The phage genera *Uliginvirus*, *Nickievirus*, and *Ghunavirus* have a high degree of similarity and extensive coverage with phages in the database, indicating that those genera have been previously described (Supp figure 4). Phage *Cruxvirus imai* Cruim01 shows high level of identity with the bacterium *P. syringae*, due to the presence of homologous prophages sequence in the bacterial genome. It was predicted that *Cruxvirus imai* Cruim01 has a lysogenic lifestyle (Supp table 3), which is further supported by the homology of this phage with the host genome. Indeed, a recent study showed that in around 99.3% of the genomes of roughly 1,500 *P. viridiflava* ATUE5 that were isolated from *Arabidopsis thaliana*, there were on average 2 prophage sequences per bacterial genome [52]. According to another study, which analysed 13,713 bacterial genomes, the average number of prophages per genome is 3.24.[53]. Finally, a recent study has shown that deleterious prophages in the *P. syringae* genome, which are incomplete phage genomes, can still be utilised by the bacterium in the form of tailocins to kill sensitive strains [54].

**Table 3:**
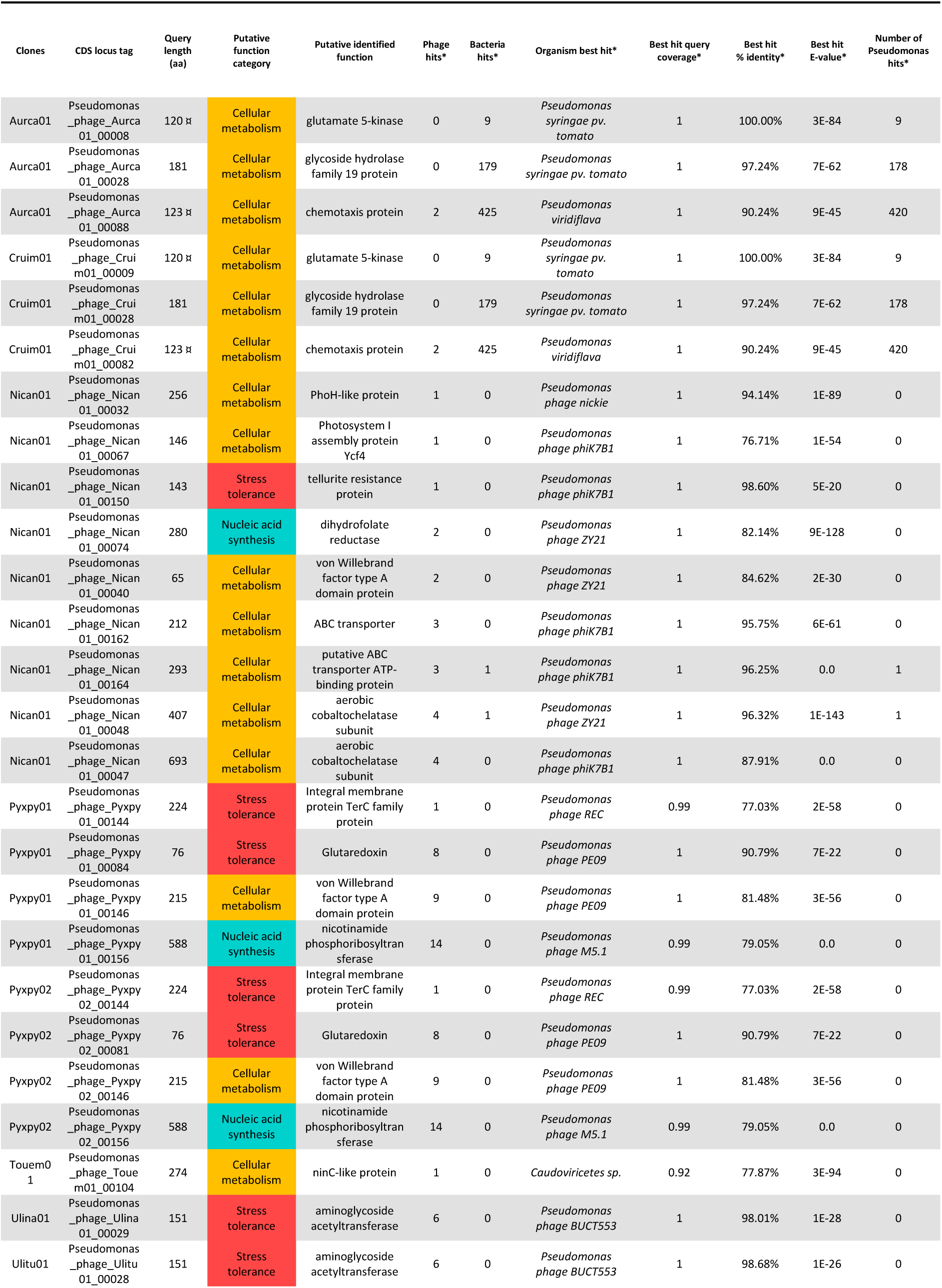
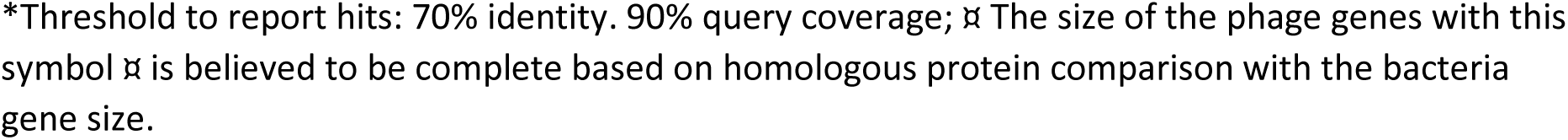
Putative auxiliary metabolic genes (AMGs) found in the *P. syringae* phage genomes classified into functional categories and comparative genomic analysis.

| Clones | CDS locus tag | Query length (aa) | Putative function category | Putative identified function | Phage hits* | Bacteria hits* | Organism best hit* | Best hit query coverage* | Best hit % identity* | Best hit E-value* | Number of Pseudomonas hits* |
| --- | --- | --- | --- | --- | --- | --- | --- | --- | --- | --- | --- |
| Aurca01 | Pseudomonas _phage_Aurca 01_00008 | 120 | Cellular metabolism | glutamate 5-kinase | 0 | 9 | *Pseudomonas syringae* pv. *tomato* | 1 | 100.00% | 3E-84 | 9 |
| Aurca01 | Pseudomonas _phage_Aurca 01_00028 | 181 | Cellular metabolism | glycoside hydrolase family 19 protein | 0 | 179 | *Pseudomonas syringae* pv. *tomato* | 1 | 97.24% | 7E-62 | 178 |
| Aurca01 | Pseudomonas _phage_Aurca 01_00088 | 123 | Cellular metabolism | chemotaxis protein | 2 | 425 | *Pseudomonas viridiflava* | 1 | 90.24% | 9E-45 | 420 |
| Cruim01 | Pseudomonas _phage_Cruim01_00009 | 120 | Cellular metabolism | glutamate 5-kinase | 0 | 9 | *Pseudomonas syringae* pv. *tomato* | 1 | 100.00% | 3E-84 | 9 |
| Cruim01 | Pseudomonas _phage_Cruim01_00028 | 181 | Cellular metabolism | glycoside hydrolase family 19 protein | 0 | 179 | *Pseudomonas syringae* pv. *tomato* | 1 | 97.24% | 7E-62 | 178 |
| Cruim01 | Pseudomonas _phage_Cruim01_00082 | 123 | Cellular metabolism | chemotaxis protein | 2 | 425 | *Pseudomonas viridiflava* | 1 | 90.24% | 9E-45 | 420 |
| Nican01 | Pseudomonas _phage_Nican 01_00032 | 256 | Cellular metabolism | PhoH-like protein | 1 | 0 | *Pseudomonas phage nickie* | 1 | 94.14% | 1E-89 | 0 |
| Nican01 | Pseudomonas _phage_Nican 01_00067 | 146 | Cellular metabolism | Photosystem I assembly protein Ycf4 | 1 | 0 | *Pseudomonas phage phiK7B1* | 1 | 76.71% | 1E-54 | 0 |
| Nican01 | Pseudomonas _phage_Nican 01_00150 | 143 | Stress tolerance | tellurite resistance protein | 1 | 0 | *Pseudomonas phage phiK7B1* | 1 | 98.60% | 5E-20 | 0 |
| Nican01 | Pseudomonas _phage_Nican 01_00074 | 280 | Nucleic acid synthesis | dihydrofolate reductase | 2 | 0 | *Pseudomonas phage ZY21* | 1 | 82.14% | 9E-128 | 0 |
| Nican01 | Pseudomonas _phage_Nican 01_00040 | 65 | Cellular metabolism | von Willebrand factor type A domain protein | 2 | 0 | *Pseudomonas phage ZY21* | 1 | 84.62% | 2E-30 | 0 |
| Nican01 | Pseudomonas _phage_Nican 01_00162 | 212 | Cellular metabolism | ABC transporter | 3 | 0 | *Pseudomonas phage phiK7B1* | 1 | 95.75% | 6E-61 | 0 |
| Nican01 | Pseudomonas _phage_Nican 01_00164 | 293 | Cellular metabolism | putative ABC transporter ATP-binding protein | 3 | 1 | *Pseudomonas phage phiK7B1* | 1 | 96.25% | 0.0 | 1 |
| Nican01 | Pseudomonas _phage_Nican 01_00048 | 407 | Cellular metabolism | aerobic cobaltochelatase subunit | 4 | 1 | *Pseudomonas phage ZY21* | 1 | 96.32% | 1E-143 | 1 |
| Nican01 | Pseudomonas _phage_Nican 01_00047 | 693 | Cellular metabolism | aerobic cobaltochelatase subunit | 4 | 0 | *Pseudomonas phage phiK7B1* | 1 | 87.91% | 0.0 | 0 |
| Pyxpy01 | Pseudomonas _phage_Pyxpy 01_00144 | 224 | Stress tolerance | Integral membrane protein TerC family protein | 1 | 0 | *Pseudomonas phage REC* | 0.99 | 77.03% | 2E-58 | 0 |
| Pyxpy01 | Pseudomonas _phage_Pyxpy 01_00084 | 76 | Stress tolerance | Glutaredoxin | 8 | 0 | *Pseudomonas phage PE09* | 1 | 90.79% | 7E-22 | 0 |
| Pyxpy01 | Pseudomonas _phage_Pyxpy 01_00146 | 215 | Cellular metabolism | von Willebrand factor type A domain protein | 9 | 0 | *Pseudomonas phage PE09* | 1 | 81.48% | 3E-56 | 0 |
| Pyxpy01 | Pseudomonas _phage_Pyxpy 01_00156 | 588 | Nucleic acid synthesis | nicotinamide phosphoribosyltransferase | 14 | 0 | *Pseudomonas phage M5.1* | 0.99 | 79.05% | 0.0 | 0 |
| Pyxpy02 | Pseudomonas _phage_Pyxpy 02_00144 | 224 | Stress tolerance | Integral membrane protein TerC family protein | 1 | 0 | *Pseudomonas phage REC* | 0.99 | 77.03% | 2E-58 | 0 |
| Pyxpy02 | Pseudomonas _phage_Pyxpy 02_00081 | 76 | Stress tolerance | Glutaredoxin | 8 | 0 | *Pseudomonas phage PE09* | 1 | 90.79% | 7E-22 | 0 |
| Pyxpy02 | Pseudomonas _phage_Pyxpy 02_00146 | 215 | Cellular metabolism | von Willebrand factor type A domain protein | 9 | 0 | *Pseudomonas phage PE09* | 1 | 81.48% | 3E-56 | 0 |
| Pyxpy02 | Pseudomonas _phage_Pyxpy 02_00156 | 588 | Nucleic acid synthesis | nicotinamide phosphoribosyltransferase | 14 | 0 | *Pseudomonas phage M5.1* | 0.99 | 79.05% | 0.0 | 0 |
| Touem01 | Pseudomonas _phage_Touem01_00104 | 274 | Cellular metabolism | ninC-like protein | 1 | 0 | *Caudoviricetes* sp. | 0.92 | 77.87% | 3E-94 | 0 |
| Ulina01 | Pseudomonas _phage_Ulina 01_00029 | 151 | Stress tolerance | aminoglycoside acetyltransferase | 6 | 0 | *Pseudomonas phage BUCT553* | 1 | 98.01% | 1E-28 | 0 |
| Ulitu01 | Pseudomonas _phage_Ulitu 01_00028 | 151 | Stress tolerance | aminoglycoside acetyltransferase | 6 | 0 | *Pseudomonas phage BUCT553* | 1 | 98.68% | 1E-26 | 0 |

\*Threshold to report hits: 70% identity. 90% query coverage; ⌘ The size of the phage genes with this symbol ⌘ is believed to be complete based on homologous protein comparison with the bacteria gene size.

This phage collection has an interesting genomic characteristic related to packaging systems of their DNA in their capsids (table 2). The genomes of our collection have two types of packaging mechanisms: direct terminal repeats (DTR) present in 32% of the phages and headful without a pac site present in 20%. These mechanisms are used by double-stranded DNA phages to circularise their genomic DNA after injection in the host bacterial cell [37]. In contrast, in 28% of the phages, no termini system was identified. In these cases, the terminase gene was conventionally used as the genome origin [55]. In addition, the Lyrsu02 clones appears to have an unusual 5’ COS-like motif with 190 base pairs cohesive sequences (Supp figure 5). It is worth noting that no known phage with cohesive sequences longer than 20 bp has been described so far. The Lyrsu01 clone also showed unusual packaging features, with the potential presence of two very distinct termini in the same genome: a 190 bp 5’ COS motif like Lyrsu02 and a 3’ COS motif (Supp figure 5). Further experimental investigation is required to confirm the identification of these potential new termini in this *Lyravirus sulafat* species.

### 5.1.4 *P. syringae* phages have consistent gene architecture within each genus and they possess various auxiliary metabolic genes (AMGs)

The phage genomes in our collection were compared based on functional annotations and their assignment to functional modules, including DNA metabolism, host lysis, lysogeny, potential auxiliary metabolic genes (AMGs), and structure, packaging, and assembly (refer to Supp table 4). After annotation, an average of 48.2% of the proteins have been annotated with a putative known function. In general, functional annotations for *P. syringae* phages are limited due to inadequate and poor databases, which requires further study [45].

The gene architecture and synteny of phages is conserved within the same genus, but each genus has a specific diversity of architectures (figure 3). Furthermore, the six genera with a single phage clone that cannot be compared were not further detailed in this section.

The four phages in the *Orionvirus* genus share at least 95% inter-genomic similarity in the *saiph* species and 94% between *mintaka* and *saiph* species. The primary distinction between these two species is the existence of an endonuclease (Pseudomonas_phage_Orimi01_00054) and a putative ERF family protein (Pseudomonas_phage_Orimi01_00055) in *Orionvirus mintaka*, which are associated with DNA metabolism and are not present in the other species. One notable characteristic of this genus in the host lysis functional module is the presence of a lysozyme that catalyses the destruction of the bacterial cell wall, followed by the o-spanin protein that disrupts the outer membrane. In the Orisa03 phage, this protein is divided into two parts: the inner membrane spanin component (Pseudomonas_phage_Orisa03_00034) and the outer membrane spanin component (Pseudomonas_phage_Orisa03_00035) [56].

The *Dracovirus* genus comprises four distinct phages across two species: *altais*, which is composed of nearly identical clones (99.9% similarity), and *eltanin*, which shares 94.8% similarity with the *altais* species. This genus of phage has two interesting distinctive features: the presence of two proteins involved in DNA metabolism (HNH homing endonucleases) and a host lysis module composed of a lysozyme and a putative Rz-like protein involved in the final step of bacterial lysis [57]. *Toucanavirus emiw* also possesses the HNH homing endonuclease protein. One difference between the two species is that *Dracovirus eltanin* lacks a block of four consecutive genes, consisting of three hypothetical proteins followed by a putative acetyltransferase of unknown function.

The *Lyravirus sulafat* species is composed of three distinct phages (with 99% similarity) that share most of their genes, with only a few differences in hypothetical proteins. The same applies to the *Pyxisvirus pyxidis* species (99% identity) which consists of two very similar phages. A notable difference in this genus is the high number of tRNAs in phage Pyxpy01, which has 19 compared to 6 in Pyxpy02 (table 2). This could explain the 1732 nucleotide difference in genome size for Pyxpy01. This genus has a higher abundance of tRNA compared to the average of 1.6 tRNAs per genome in the collection. Phages use their own DNA and RNA polymerase, but rely on the host cell for protein synthesis. It is surprising to find tRNAs in phage genomes, but their roles have been found to improve the synthesis of viral proteins [58]. Our dataset also shows that temperate phages have fewer tRNAs than virulent phages, likely due to their GC content similar to the host [51].

Additionally, the genus *Ghunavirus* currently has 10 species listed on the ICTV (as of Jan 2024), with two new species being described in this study. All species in this genus share the functional module of structure, packaging, and assembly, as well as the host lysis module (type II holin and an Rz-like lysis protein). However, there are significant nucleotide differences between species in the tail fibre protein, which could indicate a different range of phage hosts within this genus [59].

Certain phages exhibit similar gene architectures despite belonging to different genera. For instance, the genera *Aurigavirus* and *Cruxvirus* share a gene architecture (figure 3) and have an intergenomic similarity of 68.90%. Similarly, the genera *Ghunavirus* and *Lepusvirus* share a gene architecture and have an intergenomic similarity of 58.72%. The high genetic homology between the phage genera *Ghunavirus* and *Lepusvirus*, and the phage genera *Aurigavirus* and *Cruxvirus*, may indicate a speciation event. It is plausible that these two pairs of genera were a single genus before, and speciation driven by an active host-virus dynamic has allowed them to diversify [60]. This argument is suggested in particular because they share the bacterial host of isolation (CC1557 from PG10b for the *Aurigavirus* and *Cruxvirus* genera; CC0663 from PG07 for the *Ghunavirus* and *Lepusvirus* genera). Research conducted by de Leeuw *et al.* demonstrated that a phage isolated from a particular strain can enhance its predation efficiency of a different bacterial strain by selecting mutations in the tail-related region [61].

The prediction of the phage lifestyle aligns with the functions of the associated genes found in their genomes (Supp table 3). Out of the 25 phages, 22 have a predicted lytic lifestyle, while the remaining three are predicted as temperate phage, each containing at least one integrase. *Cruxvirus imai* Cruim01 also has a CII repressor, *Toucanavirus emiw* Touem01 has a ParA-like partition protein, and *Aurigavirus capella* Aurca01 has a CI repressor, a Cro-like protein, and a putative transposase [62].

However, the validation of their lysogenic lifestyle still needs experimental verification of integration in the host genome or maintenance as an episome in the host cell. Additionally, only 3 out of 25 phages were predicted as temperate, which is only 12%. Isolating and purifying from turbid plaques can be more challenging than from clear ones. The low infection capacity of some phages, which results in the formation of turbid plaques, is often associated with a lysogenic lifestyle. However, plaque morphology is not only linked to the lifestyle but also to the culture medium used and the physiological state of the bacterial host [63].

The present study identifies potential AMGs within phage genomes. These genes can regulate host cell metabolism upon infection and modulate phage replication efficiency. These genes are acquired by phages via horizontal transfer from their bacterial host [64]. Thus, the detected AMGs exhibit strong homologies with other *P. syringae* phage proteins and with the host bacterium. AMGs have been extensively studied in aquatic ecosystems, the best-known example being the cyanophages, which encode numerous genes involved in biological processes like photosynthesis, stress tolerance, nucleic acid synthesis, carbon and cellular metabolism to enhance the growth of their host [27]. Our study of the 25 *P. syringae* phage genomes revealed 26 potential AMGs, but only 4 genes have exactly the same length as the homolog in the database, raising questions about their functionality. Nevertheless, it should be noted that the threshold used to validate potential AMGs from this phage collection is stricter than in other studies [65].

The 26 potential AMGs in our *P. syringae* phages genomes can be classified into three putative functional categories: cellular metabolism (61.5%), stress tolerance (26.9%), and nucleic acid synthesis (11.5%), as presented in table 3. Three different AMGs (one is found twice) in particular are likely to be complete and have a high number of hits with *Pseudomonas* bacteria in two of our temperate phages: the *Aurigavirus capella* Aurca01 and *Cruxvirus imai* Cruim01 phages (table 3). The first one is a gene encoding the putative function of glutamate 5-kinase (Pseudomonas_phage_Aurca01_00008 and Pseudomonas_phage_Cruim01_00009). It is involved in osmotic protection as it catalyses and controls proline synthesis [66]. The second encodes a potential chemotaxis function (Pseudomonas_phage_Aurca01_00088 and Pseudomonas_phage_Cruim01_00082) that is involved in bacterial motility. This is the process by which bacteria move along gradients of attraction and can aid in their escape from non-motile phages that are diffusing [67].

Among all the phages, Nican01 has the highest number of potential AMGs, with 9 identified in its genome, belonging to all three putative functional categories. A common gene within the *Nickievirus* genus genomes encodes a photosystem I complex protein (Pseudomonas_phage_Nican01_00067), with the potential to enhance host photosynthesis and material biosynthesis (table 3). This gene is present in plants, cyanobacteria, and their associated phages [68]. For instance, it was demonstrated that phage S-PM2 encodes AMGs *psbA* and *psbD*, which are homologues of bacterial photosynthetic proteins II D1 and D2 [69]. Bacteria infected with this phage are able to maintain their photosynthetic capacity thanks to the phage’s AMGs until lysis [70]. The presence of this photosynthesis-associated gene in genomes of phages of *P. syringae* is unexpected and requires further investigation.

Interestingly, *Nickievirus* genus phages code for a tellurite resistance protein (Pseudomonas_phage_Nican01_00150) and the *Pyxisvirus* genus ones contain a gene that codes for an integral membrane protein of the *terC* family (Pseudomonas_phage_Pyxpy01_00144 and Pseudomonas_phage_Pyxpy02_00144 with a size of 224 aa) (table 3). The *terC* gene (whole *terC* gene = 346 aa) is one of seven genes that comprise the bacterial tellurite resistance determinant in plasmid R478 (*terZABCDEF*) [71]. This plasmid is present in several bacteria of the Enterobacteriaceae family and in the opportunistic bacterium *Serratia marcescens*, from which the plasmid was originally isolated. The *tmp* gene (218 aa) is the bacterial tellurite resistance determinant in the *P. syringae* genome, but it is not present in the genomes of *P. syringae* phages. According to Peng *et al.*, those genes are classified as stress-tolerance due to the toxicity of tellurite, a metalloid, to bacteria among other organisms [72].

It is important to remind that the presence of incomplete genes involved in tellurite resistance in phage genomes does not necessarily indicate that phages can provide this resistance. Indeed, the function of these genes in bacterial genomes, and particularly in phage genomes, is not yet fully understood [71]. Nevertheless, one hypothesis suggests that these potential AMGs present in phage genomes may increase or maintain tellurite resistance in infected bacteria. It has been advocated that a high abundance of AMGs would be selected in soils stressed by organochlorine pesticide pollution [65], as they would increase host fitness. This could imply that the soil of apricot trees presenting symptoms of bacterial canker constitutes a stressful environment for the bacteria due to the high proportion of AMGs found in the phage genome.

In summary, 8 out of 25 phages (32%) have potential AMGs present in their genome. It should be noted that there is currently a lack of a database and pipeline to annotate and identify AMGs in a robust and reliable manner [28]. Future research could aim to validate the function of these particular genes in the laboratory. Additionally, the stringent threshold used here may have excluded other AMGs. Also, even when proteins have low homology, their function can be conserved in phages. For instance, it has been demonstrated that the photosystem I sequence carried by phage PsaA differs from the photosystem I sequences carried by its cyanobacterial hosts (*Prochlorococcus* and *Synechococcus*), possibly due to recombination between the two hosts’ sequences [68]. This threshold could be extended by including the presence of specific protein domains, which function may be conserved despite divergence from the rest of the protein.

## 6. Conclusion

In spite of the renewed interest in phages as biocontrol agents against plant disease, the relative paucity of genetic and phenotypic data for phages of plant pathogenic bacteria makes it difficult to estimate their biogeographical distribution and potential as biocontrol agents. The main objective of this study was to isolate a diverse range of phages from *P. syringa*e and characterise their genome. It was hypothesised that increasing the number of enrichment strains would result in an extensive phage diversity. The results indicate that a wide genetic diversity of phages was successfully isolated. We have identified 25 new *P. syringae* phages, which belong to 18 new species in 14 genera, 11 of which are new. These genera are highly distinct from each other, with almost no inter-genomic homology (with the exception of 2 pairs of genera), indicating that the gene architectures and content are also highly specific to each genus. Three phage genera have been predicted to be temperate and possess genes that support this assumption. A discovery that requires further exploration and verification is the potentially new DNA packaging systems identified in the *Lyravirus sulafat* species. Finally, this study has highlighted diverse numerous potential AMGs, including genes involved in bacterial tellurite resistance, a toxic metalloid for organisms, which was previously found in the genome of phages from *P. syringae pv. actinidae*. The presence of these genes in the phage genome raises questions about their role in microbial metabolism and their implication in stress conditions in the host environment. This study increases the INPHARED databases of *P. syringae* phages by 23%, which remains relatively scarce compared to phages from *E. coli* and other more extensively studied bacterial species. Our genetic description of the phages provides a basis for future research into this host-virus system in agricultural ecosystems. Furthermore, a more comprehensive understanding of the genomics of *P. syringae* phages would allow a better assessment of their impact on the diversification and dynamics of the host population.

## Acknowledgements

We are grateful to I. Bornard head of the PACA centre’s 3A microscopy platform for performing the TEM phage analysis. We also thank the UERI team in Gotheron for helping us to collect samples in the apricot orchards. We would like to express our gratitude to C. Persyn and H. Batina for their valuable advice on DNA extraction. Finally, we are thankful to G.H. Haustant, L. Ma. Biomics Platform, C2RT, Institut Pasteur, Paris, France, supported by France Génomique (ANR-10-INBS-09) and IBISA.

