## Supplementary material for "Novel phages of *Pseudomonas syringae* unveil numerous potential auxiliary metabolic genes"

[*] Corresponding authors

Supp Table 1 to 3

Supp Figure 1 to 5

Supp Table 1: Characteristics of the orchards used to sample the phage collection.

| **Orchard name** | **Variety** | **Grafting** | **Importance of symptoms in the orchard** | **Treatments used** |
|---|---|---|---|---|
| Hybrid 2015/2016 | Hybrid | No grafting | High | Against ACL* |
| CaPReD | Anégat / Frisson | Peach tree (Monclar) at 120 cm | Medium | Without treatment |
| Core Collection | 150 varieties | Apricot tree (Manicot) at 60 cm | Depends on variety | Without treatment |

*ACL = Apricot chlorotic leafroll caused by a phytoplasm treated by insecticides.

Supp Table 2: Phages referenced in GenBank that are the most closely related to the phages of *P. syringae* in our collection.

| Genera | Species | Clones | Clostest hits | Percent identity | Query Coverage | GenBank reference |
|---|---|---|---|---|---|---|
| *Aravirus* | *cervantes* | Arace01 | Pseudomonas phage Zuri | 84.33% | 0.14 | NC_049456.2 |
| *Aurigavirus* | *capella* | Aurca01 | Pseudomonas phage Medea1 | 96.60% | 0.35 | MW862109.1 |
| *Cruxvirus* | *imai* | Cruim01 | *Pseudomonas sy* | 92.82% | 0.27 | CP047072.1 |
| *Cygnusvirus* | *sadr* | Cygsa01 | Pseudomonas phage nickie | 72.82% | 0 | NC_042091.1 |
| *Dracovirus* | *altais* | Draal01 | Pseudomonas phage Misse | 76.57% | 0.34 | MW286267.1 |
| *Dracovirus* | *altais* | Draal02 | Pseudomonas phage Bertil | 80.21% | 0.37 | MW286266.1 |
| *Dracovirus* | *altais* | Draal03 | Pseudomonas phage Misse | 76.57% | 0.3 | MW286267.1 |
| *Dracovirus* | *eltanin* | Drael01 | Pseudomonas phage Misse | 76.57% | 0.35 | MW286267.1 |
| *Ghunavirus* | *alcor* | Ghual01 | Pseudomonas phage MR1 | 93.46% | 0.91 | MT104465.1 |
| *Ghunavirus* | *chalawan* | Ghuch01 | Pseudomonas phage MR1 | 94.57% | 0.9 | MT104465.1 |
| *Lepusvirus* | *nihal* | Lepni01 | Pseudomonas phage Henninger | 79.29% | 0.47 | NC_047922.1 |
| *Lyravirus* | *sulafat* | Lyrsu01 | Pseudomonas phage UFV-P2 | 73.41% | 0.01 | NC_018850.2 |
| *Lyravirus* | *sulafat* | Lyrsu02 | Pseudomonas phage UFV-P2 | 73.41% | 0.01 | NC_018850.2 |
| *Lyravirus* | *sulafat* | Lyrsu03 | Pseudomonas phage UFV-P2 | 73.41% | 0.01 | NC_018850.2 |
| *Nickievirus* | *ankaa* | Nican01 | Pseudomonas phage phiK7B1 | 87.68% | 0.89 | MT354569.1 |
| *Orionvirus* | *mintaka* | Orimi01 | Pseudomonas phage MR15 | 92.64% | 0.02 | MT104475.1 |
| *Orionvirus* | *saiph* | Orisa01 | Pseudomonas phage MR15 | 92.64% | 0.02 | MT104475.1 |
| *Orionvirus* | *saiph* | Orisa02 | Pseudomonas phage MR15 | 92.64% | 0.02 | MT104475.1 |
| *Orionvirus* | *saiph* | Orisa03 | Pseudomonas phage MR14 | 95.83% | 0 | MT104474.1 |
| *Pavovirus* | *peacock* | Pavpe01 | Escherichia phage Halfdan | 75.87% | 0.78 | MH362766.1 |
| *Pyxisvirus* | *pyxidis* | Pyxpy01 | Pseudomonas phage PN09 | 80.31% | 0.42 | MW175491.1 |
| *Pyxisvirus* | *pyxidis* | Pyxpy02 | Pseudomonas phage PN09 | 80.31% | 0.42 | MW175491.1 |
| *Toucanavirus* | *emiw* | Touem01 | Pseudomonas phage Dolphis | 78.48% | 0.08 | MT711888.1 |
| *Uliginvirus* | *naos* | Ulina01 | Pseudomonas phage BUCT553 | 94.31% | 0.97 | MT941682.1 |
| *Uliginvirus* | *tureis* | Ulitu01 | Pseudomonas phage BUCT553 | 94.82% | 0.97 | MT941682.1 |

For each phage in the collection, their proximity to a reference phage is estimated by the percent identity and the percent of query coverage. GenBank references are those of phages close to the collection.

Supp Table 3: Predicted lifestyle of *P. syringae* phages and identified gene function related to the lysogenic cycle.

| Genera | Species | Clones | Prediction accuracy (%) | Predicted lifestyle | Genes related to lysogenic cycle |
|---|---|---|---|---|---|
| *Aravirus* | *cervantes* | Arace01 | 87.86 | Virulent | - |
| ***Aurigavirus*** | ***capella*** | **Aurca01** | **76.09** | **Temperate** | **Integrase, repressor protein CI, Cro-like protein, putative transposase** |
| ***Cruxvirus*** | ***imai*** | **Cruim01** | **71.77** | **Temperate** | **Integrase, repressor protein CII** |
| *Cygnusvirus* | *sadr* | Cygsa01 | 96.44 | Virulent | - |
| *Dracovirus* | *altais* | Draal01 | 99.71 | Virulent | - |
| *Dracovirus* | *altais* | Draal02 | 100 | Virulent | - |
| *Dracovirus* | *altais* | Draal03 | 100 | Virulent | - |
| *Dracovirus* | *eltanin* | Drael01 | 99.7 | Virulent | - |
| *Ghunavirus* | *alcor* | Ghual01 | 97.75 | Virulent | - |
| *Ghunavirus* | *chalawan* | Ghuch01 | 98.98 | Virulent | - |
| *Lepusvirus* | *nihal* | Lepni01 | 97.83 | Virulent | - |
| *Lyravirus* | *sulafat* | Lyrsu01 | 100 | Virulent | - |
| *Lyravirus* | *sulafat* | Lyrsu02 | 100 | Virulent | - |
| *Lyravirus* | *sulafat* | Lyrsu03 | 100 | Virulent | - |
| *Nickievirus* | *ankaa* | Nican01 | 83.26 | Virulent | - |
| *Orionvirus* | *mintaka* | Orimi01 | 80.02 | Virulent | - |
| *Orionvirus* | *saiph* | Orisa01 | 80.79 | Virulent | - |
| *Orionvirus* | *saiph* | Orisa02 | 83.42 | Virulent | - |
| *Orionvirus* | *saiph* | Orisa03 | 80.99 | Virulent | - |
| *Pavovirus* | *peacock* | Pavpe01 | 99.1 | Virulent | - |
| *Pyxisvirus* | *pyxidis* | Pyxpy01 | 96.03 | Virulent | - |
| *Pyxisvirus* | *pyxidis* | Pyxpy02 | 95.04 | Virulent | - |
| ***Toucanavirus*** | ***emiw*** | **Touem01** | **89.32** | **Temperate** | **Integrase, ParA-like protein** |
| *Uliginvirus* | *naos* | Ulina01 | 99.5 | Virulent | - |
| *Uliginvirus* | *tureis* | Ulitu01 | 99.55 | Virulent | - |

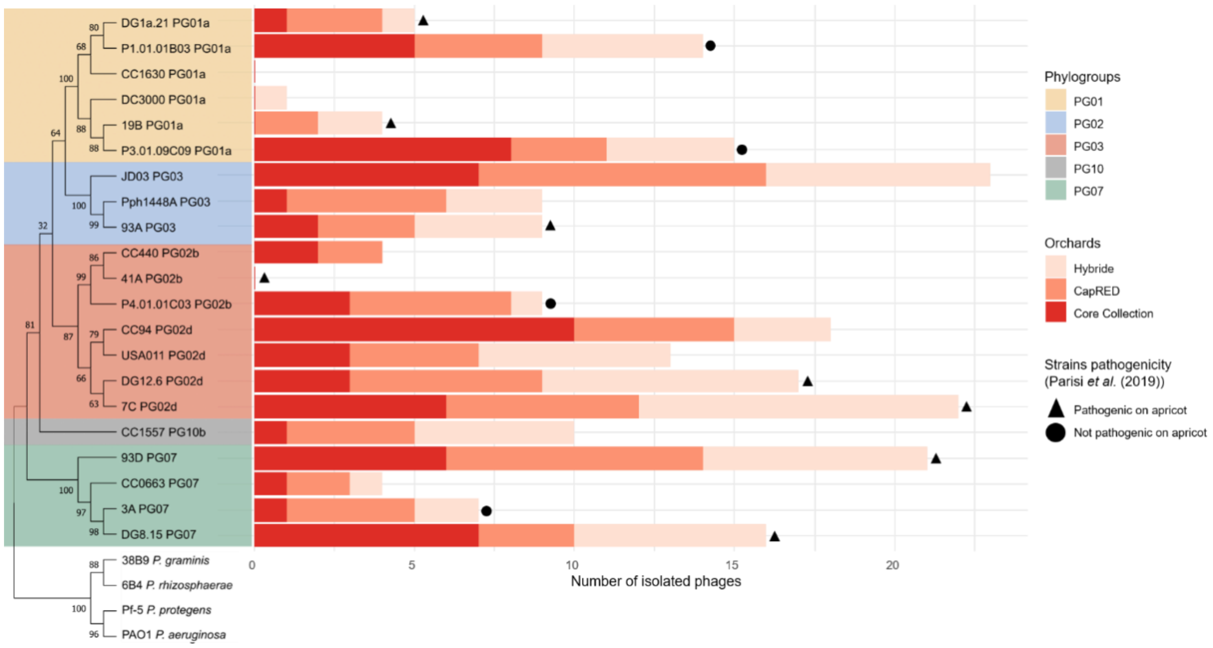


Supp Figure 1: Origin of the 221 phage isolates according to the *P. syringae* strains and the three orchards. The P. syringae strains are represented in a Neighbor Joining tree from the housekeeping cts (citrate synthase) gene and rooted with strains outside of the species complex. The four phylogroups in this tree are delimited by different coloured backgrounds (yellow: PG01a, blue: PG03, pink: PG02b and d, grey: PG10b, green: PG07). The colours of the bars represent the three orchards. The black triangles show the strains pathogenic on apricot; and the black circles indicate the strains that were not pathogenic on apricot (Parisi *et al.* 2019). Strains not marked with a triangle or circle were not tested for pathogenicity on apricot.

| Genera | Species | Clones | Orimi01 | Orisa03 | Orisa02 | Orisa01 | Cruim01 | Aurca01 | Cygsa01 | Pseudomonas phage nickie | Nican01 | Pseudomonas phage phiK7B1 | Lepni01 | Pseudomonas phage Henninger | Ghual01 | Pseudomonas phage MR1 | Ghuch01 | Lyrsu03 | Lyrsu01 | Lyrsu02 | Arace01 | Ulina01 | Ulitu01 | Pseudomonas phage BUCT553 | Drael01 | Draal01 | Draal03 | Draal02 | Pyxpy01 | Pyxpy02 | Pavpe01 | Touem01 |
|---|---|---|---|---|---|---|---|---|---|---|---|---|---|---|---|---|---|---|---|---|---|---|---|---|---|---|---|---|---|---|---|---|
| *Orionvirus* | *mintaka* | Orimi01 | 100,0 | | | | | | | | | | | | | | | | | | | | | | | | | | | | | |
| | *saiph* | Orisa03 | 91,0 | 100,0 | | | | | | | | | | | | | | | | | | | | | | | | | | | | |
| | | Orisa02 | 94,7 | 95,5 | 100,0 | | | | | | | | | | | | | | | | | | | | | | | | | | | |
| | | Orisa01 | 95,0 | 95,9 | 99,6 | 100,0 | | | | | | | | | | | | | | | | | | | | | | | | | | |
| *Cruxvirus* | *imai* | Cruim01 | 3,3 | 2,4 | 2,5 | 2,5 | 100,0 | | | | | | | | | | | | | | | | | | | | | | | | | |
| *Aurigavirus* | *capella* | Aurca01 | 3,0 | 2,6 | 3,0 | 3,0 | 68,9 | 100,0 | | | | | | | | | | | | | | | | | | | | | | | | |
| *Cygnusvirus* | *sadr* | Cygsa01 | 0,1 | 0,1 | 0,1 | 0,1 | 0,0 | 0,1 | 100,0 | | | | | | | | | | | | | | | | | | | | | | | |
| *Nickievirus* | - | Pseudomonas phage nickie | 0,0 | 0,0 | 0,0 | 0,0 | 0,0 | 0,1 | 1,4 | 100,0 | | | | | | | | | | | | | | | | | | | | | | |
| | *ankaa* | Nican01 | 0,0 | 0,0 | 0,0 | 0,0 | 0,0 | 0,1 | 1,2 | 70,0 | 100,0 | | | | | | | | | | | | | | | | | | | | | |
| | - | Pseudomonas phage phiK7B1 | 0,0 | 0,0 | 0,0 | 0,0 | 0,0 | 0,1 | 1,2 | 70,2 | 80,6 | 100,0 | | | | | | | | | | | | | | | | | | | | |
| *Lepusvirus* | *nihal* | Lepni01 | 0,0 | 0,0 | 0,0 | 0,0 | 0,0 | 0,0 | 0,0 | 0,0 | 0,0 | 0,0 | 100,0 | | | | | | | | | | | | | | | | | | | |
| *Ghunavirus* | - | Pseudomonas phage Henninger | 0,1 | 0,1 | 0,1 | 0,1 | 0,0 | 0,0 | 0,1 | 0,0 | 0,0 | 0,0 | 58,4 | 100,0 | | | | | | | | | | | | | | | | | | |
| | *alcor* | Ghual01 | 0,0 | 0,0 | 0,0 | 0,0 | 0,0 | 0,0 | 0,0 | 0,0 | 0,1 | 0,0 | 58,8 | 71,7 | 100,0 | | | | | | | | | | | | | | | | | |
| | - | Pseudomonas phage MR1 | 0,0 | 0,0 | 0,0 | 0,0 | 0,0 | 0,0 | 0,0 | 0,0 | 0,0 | 0,0 | 57,7 | 73,6 | 87,7 | 100,0 | | | | | | | | | | | | | | | | |
| | *chalawan* | Ghuch01 | 0,0 | 0,0 | 0,0 | 0,0 | 0,0 | 0,0 | 0,0 | 0,0 | 0,0 | 0,0 | 58,7 | 75,1 | 88,3 | 89,1 | 100,0 | | | | | | | | | | | | | | | |
| *Lyravirus* | *sulafat* | Lyrsu03 | 0,0 | 0,0 | 0,0 | 0,0 | 0,0 | 0,0 | 0,1 | 0,5 | 0,6 | 0,7 | 0,1 | 0,1 | 0,3 | 0,1 | 0,2 | 100,0 | | | | | | | | | | | | | | |
| | | Lyrsu01 | 0,0 | 0,0 | 0,0 | 0,0 | 0,0 | 0,0 | 0,1 | 0,5 | 0,6 | 0,7 | 0,1 | 0,1 | 0,3 | 0,1 | 0,2 | 99,2 | 100,0 | | | | | | | | | | | | | |
| | | Lyrsu02 | 0,0 | 0,0 | 0,0 | 0,0 | 0,0 | 0,0 | 0,1 | 0,5 | 0,6 | 0,7 | 0,1 | 0,1 | 0,3 | 0,1 | 0,2 | 99,1 | 99,9 | 100,0 | | | | | | | | | | | | |
| *Aravirus* | *cervantes* | Arace01 | 0,0 | 0,0 | 0,0 | 0,0 | 0,2 | 0,2 | 0,0 | 0,0 | 0,2 | 0,1 | 0,0 | 0,0 | 0,0 | 0,0 | 0,0 | 0,0 | 0,0 | 0,0 | 100,0 | | | | | | | | | | | |
| *Uliginvirus* | *naos* | Ulina01 | 0,0 | 0,0 | 0,0 | 0,0 | 0,0 | 0,0 | 0,0 | 0,0 | 0,0 | 0,1 | 0,4 | 0,3 | 0,6 | 0,6 | 0,6 | 0,2 | 0,2 | 0,2 | 1,1 | 100,0 | | | | | | | | | | |
| | *tureis* | Ulitu01 | 0,0 | 0,0 | 0,0 | 0,0 | 0,0 | 0,0 | 0,0 | 0,0 | 0,0 | 0,0 | 0,5 | 0,2 | 0,7 | 0,8 | 0,6 | 0,3 | 0,3 | 0,3 | 0,8 | 93,4 | 100,0 | | | | | | | | | |
| | - | Pseudomonas phage BUCT553 | 0,0 | 0,0 | 0,0 | 0,0 | 0,0 | 0,0 | 0,0 | 0,0 | 0,0 | 0,1 | 0,6 | 0,7 | 0,5 | 0,3 | 0,4 | 0,3 | 0,3 | 0,3 | 0,8 | 92,5 | 93,9 | 100,0 | | | | | | | | |
| *Dracovirus* | *eltanin* | Drael01 | 0,0 | 0,0 | 0,0 | 0,0 | 0,0 | 0,0 | 0,0 | 0,0 | 0,0 | 0,0 | 0,1 | 0,0 | 0,1 | 0,0 | 0,0 | 0,3 | 0,3 | 0,3 | 0,1 | 1,9 | 1,8 | 1,8 | 100,0 | | | | | | | |
| | *altais* | Draal01 | 0,0 | 0,0 | 0,0 | 0,0 | 0,0 | 0,0 | 0,0 | 0,0 | 0,0 | 0,0 | 0,1 | 0,0 | 0,1 | 0,0 | 0,0 | 0,0 | 0,0 | 0,0 | 0,1 | 1,8 | 1,8 | 1,8 | 94,8 | 100,0 | | | | | | |
| | | Draal03 | 0,0 | 0,0 | 0,0 | 0,0 | 0,0 | 0,0 | 0,0 | 0,0 | 0,0 | 0,0 | 0,1 | 0,0 | 0,1 | 0,0 | 0,0 | 0,0 | 0,0 | 0,0 | 0,1 | 1,8 | 1,8 | 1,8 | 94,8 | 99,9 | 100,0 | | | | | |
| | | Draal02 | 0,0 | 0,0 | 0,0 | 0,0 | 0,0 | 0,0 | 0,0 | 0,0 | 0,0 | 0,0 | 0,1 | 0,0 | 0,1 | 0,0 | 0,0 | 0,0 | 0,0 | 0,0 | 0,1 | 1,8 | 1,8 | 1,8 | 94,8 | 99,9 | 99,9 | 100,0 | | | | |
| *Pyxisvirus* | *pyxidis* | Pyxpy01 | 0,0 | 0,0 | 0,0 | 0,0 | 0,0 | 0,0 | 0,0 | 0,0 | 0,0 | 0,0 | 0,0 | 0,1 | 0,1 | 0,1 | 0,1 | 0,2 | 0,2 | 0,2 | 0,2 | 0,0 | 0,0 | 0,0 | 0,2 | 0,2 | 0,2 | 0,2 | 100,0 | | | |
| | | Pyxpy02 | 0,0 | 0,0 | 0,0 | 0,0 | 0,0 | 0,0 | 0,0 | 0,0 | 0,0 | 0,0 | 0,0 | 0,1 | 0,1 | 0,1 | 0,1 | 0,2 | 0,2 | 0,2 | 0,2 | 0,0 | 0,0 | 0,0 | 0,2 | 0,2 | 0,2 | 0,2 | 99,1 | 100,0 | | |
| *Pavovirus* | *peacock* | Pavpe01 | 0,0 | 0,0 | 0,0 | 0,0 | 0,0 | 0,0 | 0,0 | 0,0 | 0,0 | 0,0 | 0,0 | 0,1 | 0,0 | 0,0 | 0,0 | 0,0 | 0,0 | 0,0 | 0,5 | 0,2 | 0,2 | 0,2 | 0,0 | 0,0 | 0,0 | 0,0 | 0,0 | 0,0 | 100,0 | |
| *Toucanavirus* | *emiw* | Touem01 | 0,7 | 0,9 | 0,7 | 0,7 | 1,8 | 1,8 | 0,1 | 0,0 | 0,1 | 0,0 | 0,0 | 0,0 | 0,0 | 0,0 | 0,0 | 0,0 | 0,0 | 0,0 | 0,0 | 0,0 | 0,0 | 0,0 | 0,0 | 0,0 | 0,0 | 0,0 | 0,0 | 0,0 | 0,0 | 100,0 |

Supp Figure 2: Genetic distance matrix of the 25 *P. syringae* phages compared to the 5-close reference phages (in yellow), produced with VIRIDIC. The heatmap displays the percentage of inter-genome identity and phage genera clusters are indicated in green and in the first columns (species and clones are included for clarity purposes).

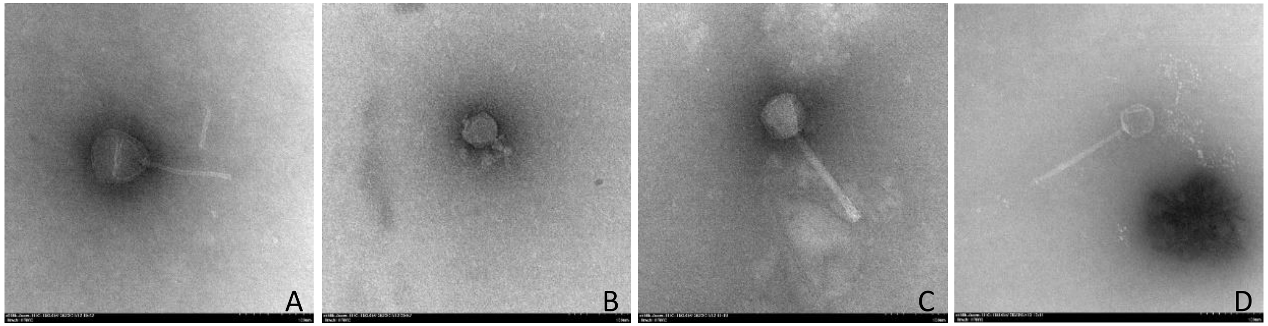


Supp Figure 3: TEM image of 4 *P. syringae* phages presenting three different morphologies: siphovirus (A, virulent phage Nican01), podovirus (B, virulent phage Lyrsu02), myovirus (C, temperate phage Touem01) and siphovirus (D, virulent phage Pavpe01).

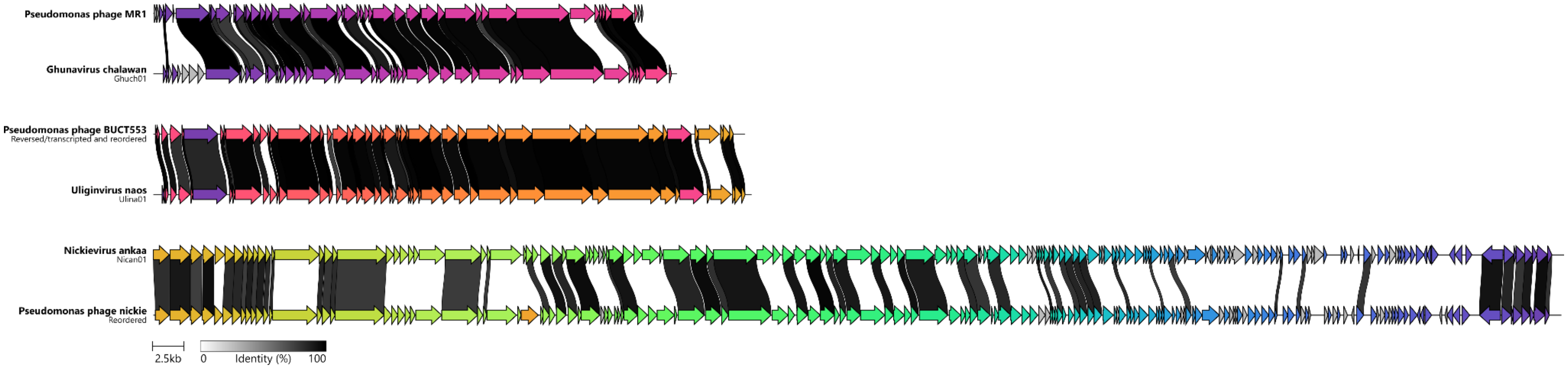


Supp Figure 4: Genomic comparison of the 3 phage genera previously described with the 3 new *P. syringae* phages belonging to them using Clinker. The genomes previously described have been reorganised or reversed to allow alignment with the new genomes. The arrows indicate the open reading frames and their direction. Link identity are presented as black to white bands and threshold was set to 80% protein identity.

Supp Figure 5: Reports on the PhageTerm analysis of the three clones of the *Lyravirus sulafat* species.

### *Lyravirus sulafat Lyrsu01 PhageTerm Analysis*

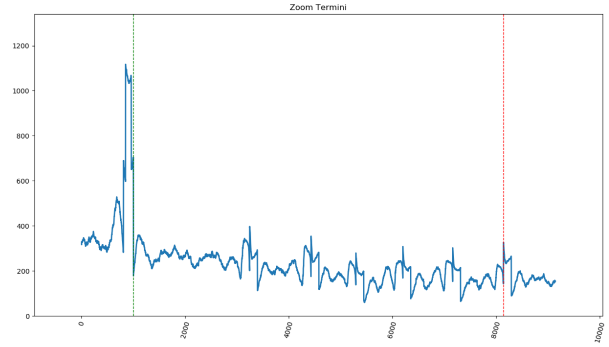


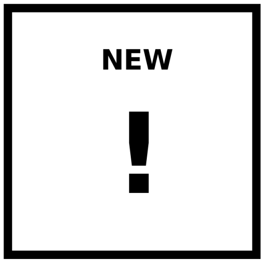


#### PhageTerm Method

| Ends | Left (red) | Right (green) | Permuted | Orientation | Class | Type |
|---|---|---|---|---|---|---|
| Redundant | 20466 | 13323 | No | NA | - | - |

| Strand | Location | T | pvalue |
|---|---|---|---|
| + | 20466 | 0.93 | 3.34e-51 |
| | 26881 | 0.92 | 2.82e-43 |
| | 39573 | 0.91 | 1.87e-51 |
| | 18525 | 0.90 | 7.09e-40 |
| | 17618 | 0.89 | 9.66e-39 |
| - | 13323 | 0.79 | 4.94e-181 |
| | 11282 | 0.26 | 5.55e-15 |
| | 6133 | 0.24 | 3.64e-04 |
| | 4405 | 0.14 | 2.12e-03 |
| | 972 | 0.12 | 9.51e-03 |

**T (Start. Pos. Cov. / Whole Cov.)**

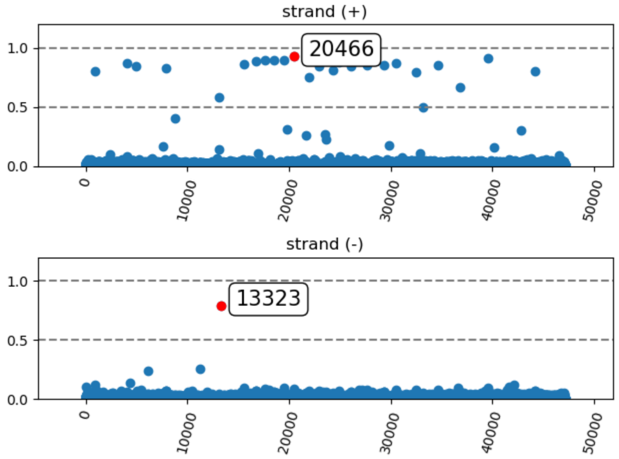


#### Li's Method

| Packaging | Termini | Forward | Reverse | Orientation |
|---|---|---|---|---|
| PAC | Fixed | Multiple-Pref. Term. | Obvious Termini | Reverse |

| Strand | Location | SPC | R |
|---|---|---|---|
| + | 13133 | 369 | 2.0 |
| | 7900 | 226 | - |
| | 4996 | 204 | - |
| | 956 | 196 | - |
| | 15568 | 193 | - |
| - | 13323 | 525 | 11.0 |
| | 11282 | 50 | - |
| | 6133 | 28 | - |
| | 4405 | 19 | - |
| | 972 | 17 | - |

**SPC**

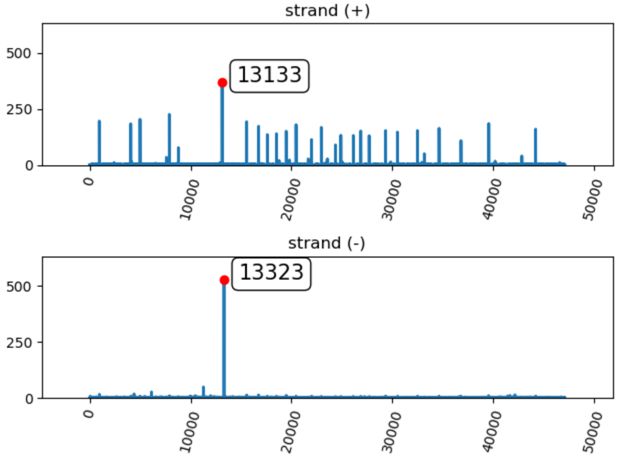

### Analysis Methodology

*PhageTerm software uses raw reads of a phage sequenced with a sequencing technology using random fragmentation and its genomic reference sequence to determine the termini position. The process starts with the alignment of NGS reads to the phage genome in order to calculate the starting position coverage (SPC), where a hit is given only to the position of the first base in a successfully aligned read (the alignment algorithm uses the lenght of the seed (default: 20) for mapping and does not accept gap or missmatch to speed up the process). Then the program apply 2 distinct scoring methods: i) a statistical approach based on the Gamma law; and ii) a method derived from LI and al. 2014 paper.*

#### General set-up and mapping informations

| | |
|---|---|
| **Phage Genome** | 47123 bp |
| **Sequencing Reads** | 70937 |
| **Mapping Reads** | 97 % |
| *OPTIONS* | |
| *Mapping Seed* | *20* |
| *Surrounding* | *20* |
| *Host Analysis* | *No* |

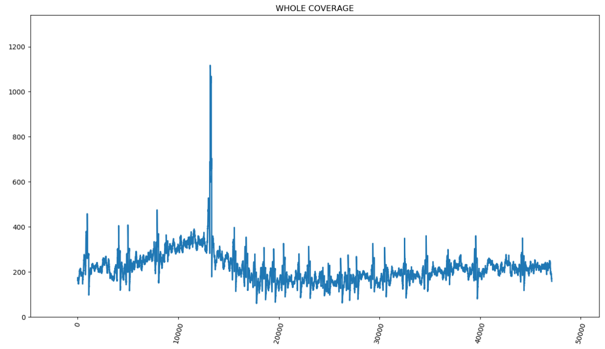


#### Highest peak of each side coverage graphics

**Whole Coverage Zoom (Left)** | **Whole Coverage Zoom (Right)**

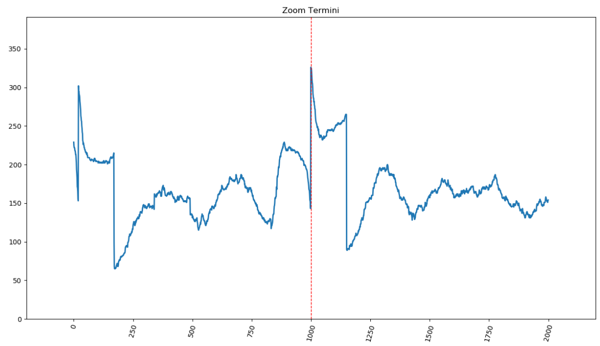


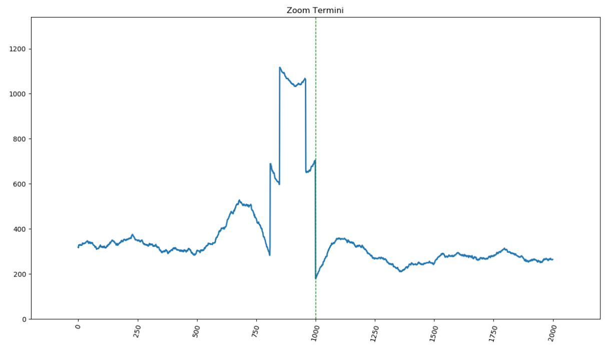


#### General controls information

| | | |
|---|---|---|
| **Whole genome coverage** | 108 | WARNING: Low (<200), Li's method could not be reliable |
| **Weak genome coverage** | 0.0 % | OK |
| **Reads lost during alignment** | 2.4 % | OK |

### i) PhageTerm method

*Reads are mapped on the reference to determine the starting position coverage (SPC) as well as the coverage (COV) in each orientation. These values are then used to compute the variable T = SPC / COV. The average value of T at positions along the genome that are not termini is expected to be 1/F, where F is the average fragment size. For the termini that depends of the packaging mode. Cos Phages: no reads should start before the terminus and therefore X=1. DTR phages: for N phages present in the sample, there should be N fragments that start at the terminus and N fragments that cover the edge of the repeat on the other side of the genome as a results T is expected to be 0.5. Pac phages: for N phages in the sample, there should be N/C fragments starting at the pac site, where C is the number of phage genome copies per concatemer. In the same sample N fragments should cover the pac site position, T is expected to be (N/C)/(N+N/C) = 1/(1+C). To assess whether the number of reads starting at a given position along the genome can be considered a significant outlier, PhageTerm first segments the genome according to coverage using a regression tree. A gamma distribution is then fitted to SPC for each segment and an adjusted p-value is computed for each position. Finally if several significant peaks are detected within a small sequence window (default: 20bp), their T values are merged.*

| | | |
|---|---|---|
| **Nearby Termini (Forward / Reverse)** | 0 / 0 | Peaks localized 20 bases around the maximum |

### ii) Li's method

*The second approach is based on the calculation and interpretation of three specific ratios R1, R2 and R3 as suggested in a previous publication from Li et al. 2014. The first ratio, is calculated as follow: the highest starting frequency found on either the forward or reverse strands is divided by the average starting frequency, R1 = (highest frequency/average frequency). Li's et al. have proposed three possible interpretation of the R1 ratio. First, if R1 < 30, the phage genome does not have any termini, and is either circular or completely permuted and terminally redundant. The second interpretation for R1 is when 30 ≤ R1 ≥ 100, suggesting the presence of preferred termini with terminal redundancy and apparition of partially circular permutations. At last if R1 > 100 that is an indication that at least one fixed termini is present with terminase recognizing a specific site. The two other ratios are R2 and R3 and the calculation is done in a similar manner. R2 is calculated using the highest two frequencies (T1-F and T2-F) found on the forward strand and R3 is calculated using the highest two frequencies (T1-R and T2-R) found on the reverse strand. To calculate these two ratios, we divide the highest frequency by the second highest frequency T2. So R2 = (T1-F / T2-F) and R3 = (T1-R / T2-R). These two ratios are used to analyze termini characteristics on each strand taken individually. Li et al. suggested two possible interpretations for R2 and R3 ratios combine to R1. When R1 < 30 and R2 < 3, we either have no obvious termini on the forward strand, or we have multiple preferred termini on the forward strand, if 30 ≤ R1 ≤ 100. If R2 > 3, it is suggested that there is an obvious unique termini on the forward strand. The same reasoning is applicable for the result of R3. Combining the results for ratios found with this approach, it is possible to make the first prediction for the viral packaging mode of the analyzed phage. A unique obvious termini present at both ends (both R2 and R3 > 3) reveals the presence of a COS mode of packaging. The headful mode of packaging PAC is concluded when we have a single obvious termini only on one strand. A whole coverage around 500X is needed for this method to be reliable.*

| | | |
|---|---|---|
| **Nearby Termini (Forward / Reverse)** | 0 / 0 | Peaks localized 20 bases around the maximum |
| **R1 - highest freq./average freq.** | 714 | At least one fixed termini is present with terminase recognizing a specific site. |
| **R2 Forw - highest freq./second freq.** | 2 | Multiple preferred termini on the forward strand. |
| **R3 Rev - highest freq./second freq.** | 11 | Unique termini on the reverse strand. |

### *Lyravirus sulafat Lyrsu02 PhageTerm Analysis*

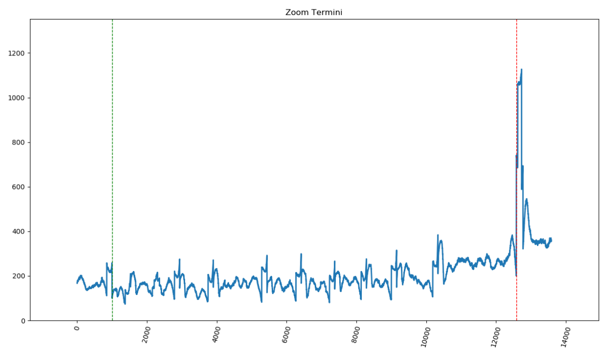


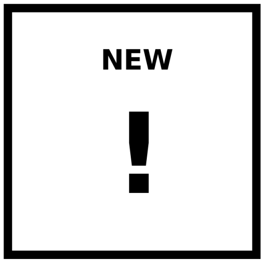


#### PhageTerm Method

| Ends | Left (red) | Right (green) | Permuted | Orientation | Class | Type |
|---|---|---|---|---|---|---|
| Redundant | 37641 | 26058 | No | NA | - | - |

| Strand | Location | T | pvalue |
|---|---|---|---|
| + | 37641 | 0.79 | 2.56e-131 |
| | 39682 | 0.29 | 1.72e-12 |
| | 44831 | 0.23 | 4.16e-07 |
| | 46559 | 0.16 | 1.82e-02 |
| | 8800 | 0.15 | 8.07e-08 |
| - | 26058 | 0.92 | 4.22e-44 |
| | 30498 | 0.91 | 4.59e-48 |
| | 34211 | 0.89 | 2.62e-62 |
| | 16324 | 0.88 | 2.71e-48 |
| | 32439 | 0.88 | 5.01e-35 |

**T (Start. Pos. Cov. / Whole Cov.)**

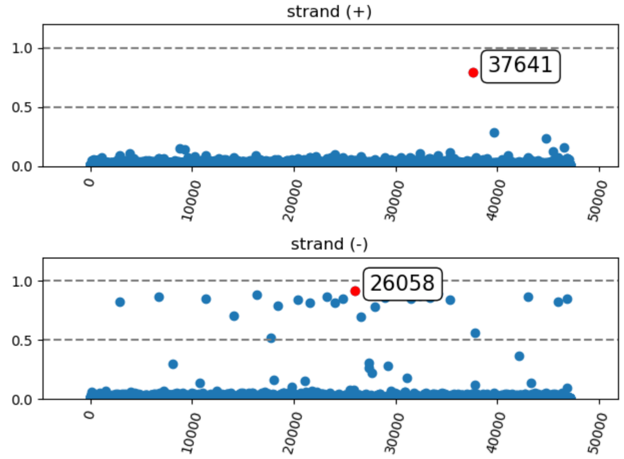


#### Li's Method

| Packaging | Termini | Forward | Reverse | Orientation |
|---|---|---|---|---|
| PAC | Fixed | Obvious Termini | Multiple-Pref. Term. | Forward |

| Strand | Location | SPC | R |
|---|---|---|---|
| + | 37641 | 541 | 9.0 |
| | 39682 | 60 | - |
| | 44831 | 36 | - |
| | 46559 | 23 | - |
| | 8800 | 21 | - |
| - | 37831 | 339 | 1.0 |
| | 43064 | 239 | - |
| | 45968 | 202 | - |
| | 6751 | 172 | - |
| | 2940 | 171 | - |

**SPC**

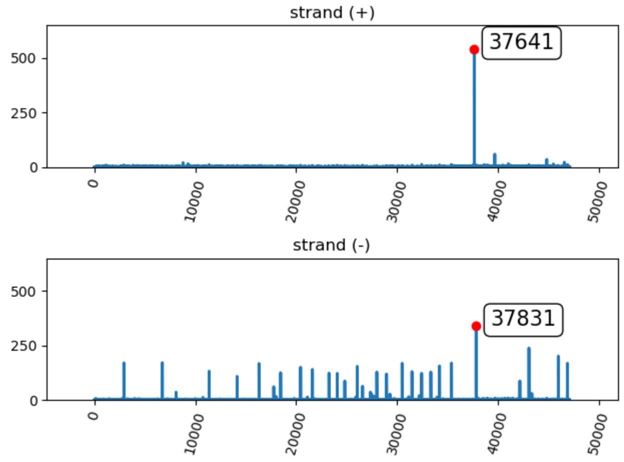

### Analysis Methodology

*PhageTerm software uses raw reads of a phage sequenced with a sequencing technology using random fragmentation and its genomic reference sequence to determine the termini position. The process starts with the alignment of NGS reads to the phage genome in order to calculate the starting position coverage (SPC), where a hit is given only to the position of the first base in a successfully aligned read (the alignment algorithm uses the lenght of the seed (default: 20) for mapping and does not accept gap or missmatch to speed up the process). Then the program apply 2 distinct scoring methods: i) a statistical approach based on the Gamma law; and ii) a method derived from LI and al. 2014 paper.*

#### General set-up and mapping informations

| | |
|---|---|
| **Phage Genome** | 47123 bp |
| **Sequencing Reads** | 71442 |
| **Mapping Reads** | 97 % |
| *OPTIONS* | |
| *Mapping Seed* | *20* |
| *Surrounding* | *20* |
| *Host Analysis* | *No* |

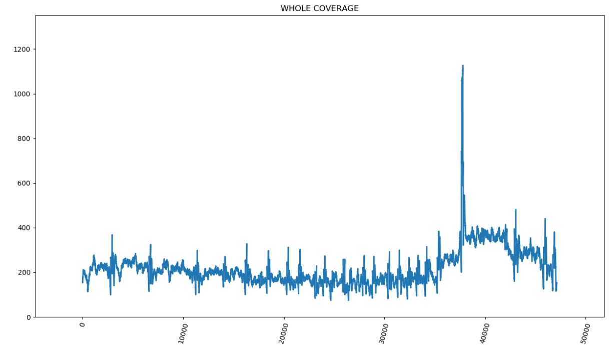


#### Highest peak of each side coverage graphics

**Whole Coverage Zoom (Left)** — **Whole Coverage Zoom (Right)**

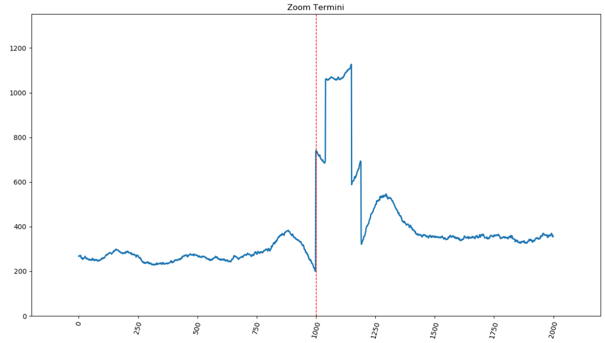


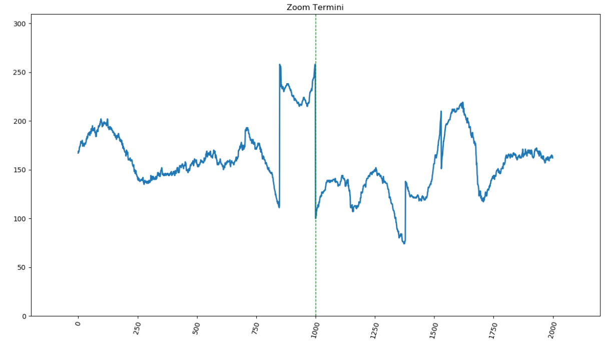


#### General controls information

| | | |
|---|---|---|
| **Whole genome coverage** | 108 | WARNING: Low (<200), Li's method could not be reliable |
| **Weak genome coverage** | 0.0 % | OK |
| **Reads lost during alignment** | 2.7 % | OK |

### i) PhageTerm method

*Reads are mapped on the reference to determine the starting position coverage (SPC) as well as the coverage (COV) in each orientation. These values are then used to compute the variable T = SPC / COV. The average value of T at positions along the genome that are not termini is expected to be 1/F, where F is the average fragment size. For the termini that depends of the packaging mode. Cos Phages: no reads should start before the terminus and therefore X=1. DTR phages: for N phages present in the sample, there should be N fragments that start at the terminus and N fragments that cover the edge of the repeat on the other side of the genome as a results T is expected to be 0.5. Pac phages: for N phages in the sample, there should be N/C fragments starting at the pac site, where C is the number of phage genome copies per concatemer. In the same sample N fragments should cover the pac site position, T is expected to be (N/C)/(N+N/C) = 1/(1+C). To assess whether the number of reads starting at a given position along the genome can be considered a significant outlier, PhageTerm first segments the genome according to coverage using a regression tree. A gamma distribution is then fitted to SPC for each segment and an adjusted p-value is computed for each position. Finally if several significant peaks are detected within a small sequence window (default: 20bp), their T values are merged.*

| | | |
|---|---|---|
| **Nearby Termini (Forward / Reverse)** | 0 / 0 | Peaks localized 20 bases around the maximum |

### ii) Li's method

*The second approach is based on the calculation and interpretation of three specific ratios R1, R2 and R3 as suggested in a previous publication from Li et al. 2014. The first ratio, is calculated as follow: the highest starting frequency found on either the forward or reverse strands is divided by the average starting frequency, R1 = (highest frequency/average frequency). Li's et al. have proposed three possible interpretation of the R1 ratio. First, if R1 < 30, the phage genome does not have any termini, and is either circular or completely permuted and terminally redundant. The second interpretation for R1 is when 30 ≤ R1 ≥ 100, suggesting the presence of preferred termini with terminal redundancy and apparition of partially circular permutations. At last if R1 > 100 that is an indication that at least one fixed termini is present with terminase recognizing a specific site. The two other ratios are R2 and R3 and the calculation is done in a similar manner. R2 is calculated using the highest two frequencies (T1-F and T2-F) found on the forward strand and R3 is calculated using the highest two frequencies (T1-R and T2-R) found on the reverse strand. To calculate these two ratios, we divide the highest frequency by the second highest frequency T2. So R2 = (T1-F / T2-F) and R3 = (T1-R / T2-R). These two ratios are used to analyze termini characteristics on each strand taken individually. Li et al. suggested two possible interpretations for R2 and R3 ratios combine to R1. When R1 < 30 and R2 < 3, we either have no obvious termini on the forward strand, or we have multiple preferred termini on the forward strand, if 30 ≤ R1 ≤ 100. If R2 > 3, it is suggested that there is an obvious unique termini on the forward strand. The same reasoning is applicable for the result of R3. Combining the results for ratios found with this approach, it is possible to make the first prediction for the viral packaging mode of the analyzed phage. A unique obvious termini present at both ends (both R2 and R3 > 3) reveals the presence of a COS mode of packaging. The headful mode of packaging PAC is concluded when we have a single obvious termini only on one strand. A whole coverage around 500X is needed for this method to be reliable.*

| | | |
|---|---|---|
| **Nearby Termini (Forward / Reverse)** | 0 / 0 | Peaks localized 20 bases around the maximum |
| **R1 - highest freq./average freq.** | 733 | At least one fixed termini is present with terminase recognizing a specific site. |
| **R2 Forw - highest freq./second freq.** | 9 | Unique termini on the forward strand. |
| **R3 Rev - highest freq./second freq.** | 1 | Multiple preferred termini on the reverse strand. |

### *Lyravirus sulafat Lyrsu03 PhageTerm Analysis*

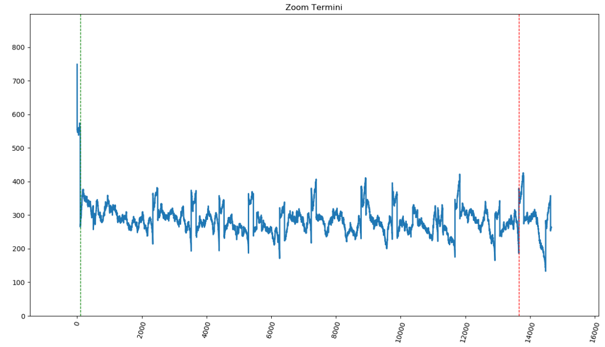


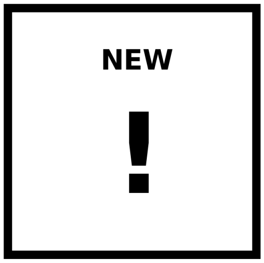


#### PhageTerm Method

| Ends | Left (red) | Right (green) | Permuted | Orientation | Class | Type |
|---|---|---|---|---|---|---|
| Redundant | 13655 | 97 | No | NA | - | - |

| Strand | Location | T | pvalue |
|---|---|---|---|
| + | 13655 | 0.83 | 3.59e-49 |
| | 38006 | 0.81 | 2.53e-48 |
| | 14484 | 0.81 | 4.00e-38 |
| | 6261 | 0.81 | 1.08e-38 |
| | 7240 | 0.80 | 2.16e-40 |
| - | 97 | 0.62 | 1.01e-119 |
| | 40058 | 0.15 | 4.12e-01 |
| | 45207 | 0.14 | 1.48e-02 |
| | 33241 | 0.13 | 1.00e+00 |
| | 33778 | 0.11 | 1.00e+00 |

**T (Start. Pos. Cov. / Whole Cov.)**

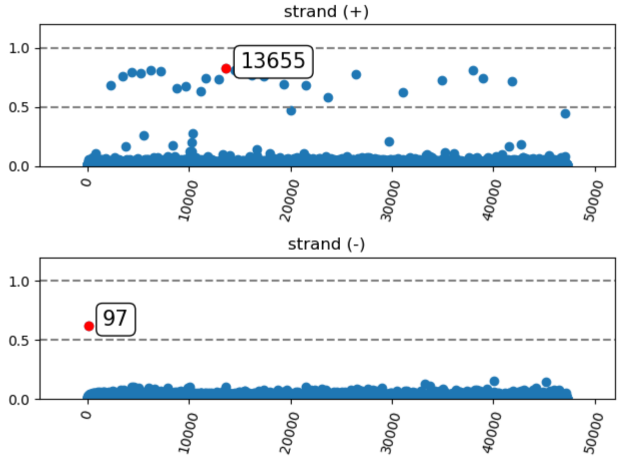


#### Li's Method

| Packaging | Termini | Forward | Reverse | Orientation |
|---|---|---|---|---|
| PAC | Fixed | Multiple-Pref. Term. | Obvious Termini | Reverse |

| Strand | Location | SPC | R |
|---|---|---|---|
| + | 47058 | 191 | 1.0 |
| | 16200 | 175 | - |
| | 13655 | 173 | - |
| | 17374 | 172 | - |
| | 38006 | 170 | - |
| - | 97 | 286 | 11.0 |
| | 45207 | 26 | - |
| | 40058 | 21 | - |
| | 4410 | 18 | - |
| | 33778 | 17 | - |

**SPC**

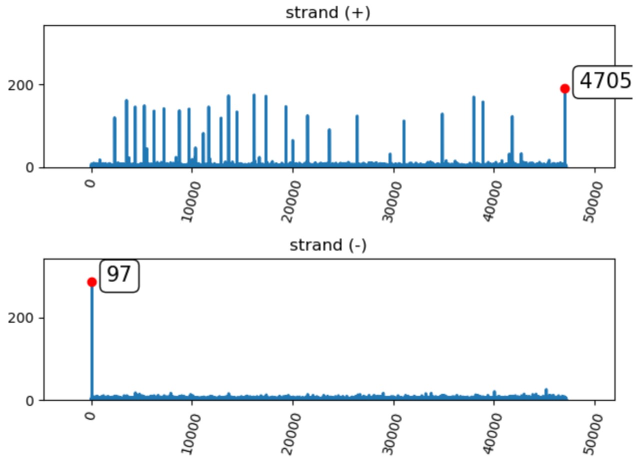

#### Analysis Methodology

*PhageTerm software uses raw reads of a phage sequenced with a sequencing technology using random fragmentation and its genomic reference sequence to determine the termini position. The process starts with the alignment of NGS reads to the phage genome in order to calculate the starting position coverage (SPC), where a hit is given only to the position of the first base in a successfully aligned read (the alignment algorithm uses the lenght of the seed (default: 20) for mapping and does not accept gap or missmatch to speed up the process). Then the program apply 2 distinct scoring methods: i) a statistical approach based on the Gamma law; and ii) a method derived from LI and al. 2014 paper.*

##### General set-up and mapping informations

| | |
|---|---|
| **Phage Genome** | 47206 bp |
| **Sequencing Reads** | 97566 |
| **Mapping Reads** | 98 % |
| *OPTIONS* | |
| *Mapping Seed* | *20* |
| *Surrounding* | *20* |
| *Host Analysis* | *No* |

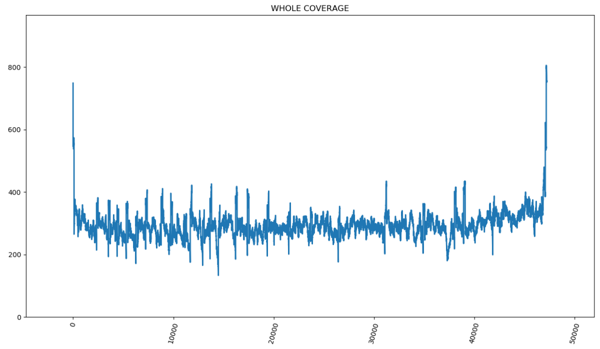


##### Highest peak of each side coverage graphics

**Whole Coverage Zoom (Left)** **Whole Coverage Zoom (Right)**

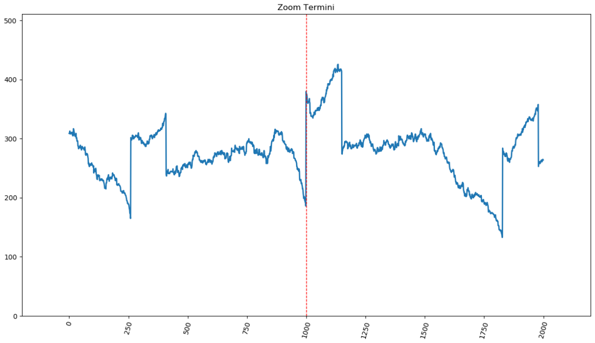


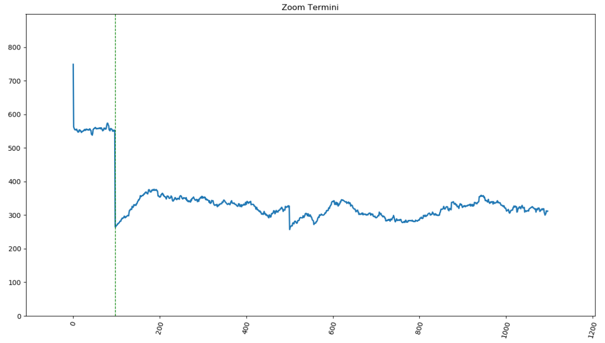


##### General controls information

| | | |
|---|---|---|
| **Whole genome coverage** | 146 | WARNING: Low (<200), Li's method could not be reliable |
| **Weak genome coverage** | 0.0 % | OK |
| **Reads lost during alignment** | 1.8 % | OK |

### i) PhageTerm method

*Reads are mapped on the reference to determine the starting position coverage (SPC) as well as the coverage (COV) in each orientation. These values are then used to compute the variable T = SPC / COV. The average value of T at positions along the genome that are not termini is expected to be 1/F, where F is the average fragment size. For the termini that depends of the packaging mode. Cos Phages: no reads should start before the terminus and therefore X=1. DTR phages: for N phages present in the sample, there should be N fragments that start at the terminus and N fragments that cover the edge of the repeat on the other side of the genome as a results T is expected to be 0.5. Pac phages: for N phages in the sample, there should be N/C fragments starting at the pac site, where C is the number of phage genome copies per concatemer. In the same sample N fragments should cover the pac site position, T is expected to be (N/C)/(N+N/C) = 1/(1+C). To assess whether the number of reads starting at a given position along the genome can be considered a significant outlier, PhageTerm first segments the genome according to coverage using a regression tree. A gamma distribution is then fitted to SPC for each segment and an adjusted p-value is computed for each position. Finally if several significant peaks are detected within a small sequence window (default: 20bp), their T values are merged.*

| | | |
|---|---|---|
| **Nearby Termini (Forward / Reverse)** | 0 / 0 | Peaks localized 20 bases around the maximum |

### ii) Li's method

*The second approach is based on the calculation and interpretation of three specific ratios R1, R2 and R3 as suggested in a previous publication from Li et al. 2014. The first ratio, is calculated as follow: the highest starting frequency found on either the forward or reverse strands is divided by the average starting frequency, R1 = (highest frequency/average frequency). Li's et al. have proposed three possible interpretation of the R1 ratio. First, if R1 < 30, the phage genome does not have any termini, and is either circular or completely permuted and terminally redundant. The second interpretation for R1 is when 30 ≤ R1 ≥ 100, suggesting the presence of preferred termini with terminal redundancy and apparition of partially circular permutations. At last if R1 > 100 that is an indication that at least one fixed termini is present with terminase recognizing a specific site. The two other ratios are R2 and R3 and the calculation is done in a similar manner. R2 is calculated using the highest two frequencies (T1-F and T2-F) found on the forward strand and R3 is calculated using the highest two frequencies (T1-R and T2-R) found on the reverse strand. To calculate these two ratios, we divide the highest frequency by the second highest frequency T2. So R2 = (T1-F / T2-F) and R3 = (T1-R / T2-R). These two ratios are used to analyze termini characteristics on each strand taken individually. Li et al. suggested two possible interpretations for R2 and R3 ratios combine to R1. When R1 < 30 and R2 < 3, we either have no obvious termini on the forward strand, or we have multiple preferred termini on the forward strand, if 30 ≤ R1 ≤ 100. If R2 > 3, it is suggested that there is an obvious unique termini on the forward strand. The same reasoning is applicable for the result of R3. Combining the results for ratios found with this approach, it is possible to make the first prediction for the viral packaging mode of the analyzed phage. A unique obvious termini present at both ends (both R2 and R3 > 3) reveals the presence of a COS mode of packaging. The headful mode of packaging PAC is concluded when we have a single obvious termini only on one strand. A whole coverage around 500X is needed for this method to be reliable.*

| | | |
|---|---|---|
| **Nearby Termini (Forward / Reverse)** | 0 / 0 | Peaks localized 20 bases around the maximum |
| **R1 - highest freq./average freq.** | 281 | At least one fixed termini is present with terminase recognizing a specific site. |
| **R2 Forw - highest freq./second freq.** | 1 | Multiple preferred termini on the forward strand. |
| **R3 Rev - highest freq./second freq.** | 11 | Unique termini on the reverse strand. |
